# Cancer-associated fibroblasts in pancreatic ductal adenocarcinoma determine response to SLC7A11 inhibition

**DOI:** 10.1101/2020.07.12.199638

**Authors:** George Sharbeen, Joshua A. McCarroll, Anouschka Akerman, Chantal Kopecky, Janet Youkhana, Jeff Holst, Cyrille Boyer, Mert Erkan, David Goldstein, Paul Timpson, Thomas R. Cox, Brooke A. Pereira, Jessica L. Chitty, Sigrid Fey, Arafath K. Najumudeen, Andrew D. Campbell, Owen J. Sansom, Rosa Mistica C. Ignacio, Stephanie Naim, Jie Liu, Nelson Russia, Julia Lee, Angela Chou, Amber Johns, Anthony Gill, Estrella Gonzales-Aloy, John Kokkinos, Val Gebski, Nigel Turner, Minoti Apte, Thomas P. Davis, Jennifer P. Morton, Koroush Haghighi, Australian Pancreatic Cancer Genome Initiative, Phoebe A. Phillips

## Abstract

Cancer-Associated Fibroblasts (CAFs) are major contributors to pancreatic ductal adenocarcinoma (PDAC) progression, through pro-tumour cross-talk and the generation of fibrosis (physical barrier to drugs). CAF inhibition is thus an ideal component of any therapeutic approach for PDAC. SLC7A11 is a cystine transporter that has been identified as a potential therapeutic target in PDAC cells. However, no prior study has evaluated the role of SLC7A11 in PDAC tumour stroma and its prognostic significance. Herein we show that high expression of SLC7A11 in PDAC tumour stroma (but not tumour cells) is independently prognostic of poorer overall survival. We demonstrate using orthogonal approaches that PDAC-derived CAFs are highly dependent on SLC7A11 for cystine uptake and glutathione synthesis, and that SLC7A11 inhibition significantly decreases their proliferation, reduces their resistance to oxidative stress and inhibits their ability to remodel collagen and support PDAC cell growth. Importantly, our paradigm-shifting work demonstrates the need to inhibit SLC7A11 in the PDAC stroma, as genetic ablation of SLC7A11 in PDAC cells alone is not enough to reduce tumour growth. Finally, our work validates that a nano-based gene-silencing drug against SLC7A11, developed by our group, reduces PDAC tumour growth, CAF activation and fibrosis in a mouse model of PDAC.

## INTRODUCTION

Pancreatic ductal adenocarcinoma (PDAC) is a lethal malignancy, with a 5-year survival rate of <9% (1). A major reason for this poor prognosis is the drug-refractory nature of PDAC caused by inherent chemoresistance mechanisms and physical barriers to drug delivery. The dense fibrotic PDAC microenvironment drives both of these mechanisms (2). Fibrosis distorts the tumour vasculature physically hindering drug access and creates a harsh hypoxic and nutrient-deprived microenvironment (2). These conditions promote the transition of cancer cells from an epithelial phenotype to a more metastatic and chemoresistant mesenchymal phenotype (2). The architects of the PDAC microenvironment are Cancer-Associated Fibroblasts (CAFs) (2, 3). CAFs are activated by signals released from PDAC cells and hypoxia, resulting in a self-perpetuating loop of excessive extracellular matrix (ECM) protein deposition that creates fibrosis (2, 3). CAFs also reciprocate pro-survival signalling to PDAC cells, thus promoting PDAC cell survival and epithelial to mesenchymal transition (2, 3). This makes stromal remodelling and inhibition of CAF activity an important consideration for PDAC therapeutic approaches.

CAFs and PDAC cells share an oxygen/nutrient poor microenvironment. PDAC cells have altered their metabolism to survive and proliferate in this stressful microenvironment (4). These alterations can lead to metabolic addictions that can be therapeutically exploited. A potential target that has gained significant interest is the X_c_^-^ amino acid antiporter, which imports cystine into the cell, in exchange for glutamate (5-7). X_c_^-^ is a heterodimer of solute carrier 3A2 (SLC3A2; membrane anchor) and solute carrier 7A11 (SLC7A11, also known as xCT; amino acid transporter) (5-7). This transporter sits at the crux of multiple metabolic activities necessary for cancer cell survival, including protein synthesis and redox regulation. First, cystine transported by SLC7A11 is reduced to cysteine, which is an irreplaceable component of proteins, that is required for disulphide bond formation. Second, cysteine is also the rate-limiting amino acid in the synthesis of the potent antioxidant glutathione (GSH) (8). GSH is important in PDAC cell survival as KRAS-driven metabolic changes, pro-tumour signalling and microenvironment-driven hypoxia increase intracellular oxidative stress (9). Without this protection, uncontrolled oxidative stress could compromise cell survival by damaging DNA and proteins.

In light of these critical roles, SLC7A11 has been identified as a prognostic factor and potential therapeutic target in a number of cancers (10-15). In PDAC, Lo et al (16) demonstrated that SLC7A11 was upregulated in PDAC cells under oxidative stress and cystine deprivation *in vitro*. They subsequently showed that SLC7A11 inhibitor sulfasalazine (SSZ) significantly reduced subcutaneous PDAC tumour growth (17). Since then, additional studies have demonstrated the therapeutic potential of inhibiting or genetically ablating SLC7A11 in PDAC cells (18-22). While these findings were promising, a key limitation of all the above studies was that they ignored the role of SLC7A11 in CAFs or the impact of CAFs on PDAC cell sensitivity to SLC7A11 inhibition. This is a critical gap in our knowledge for therapeutic inhibition of SLC7A11 in PDAC, given the prominent cross-talk between CAFs and PDAC cells and their impact on PDAC drug sensitivity. In addition, evidence suggests that amino acids can be exchanged between tumour cells and stromal cells to help overcome nutrient deficiencies and drive tumour progression (23, 24). Therefore, it is possible that SLC7A11 may play a similar role in PDAC/CAF metabolic cross-talk because of its ability to regulate both glutamate efflux and cysteine production.

We hypothesised that SLC7A11 inhibition in CAFs had the potential to directly inhibit a key cellular target in PDAC and to break a potential nutrient feeding axis between CAFs and PDAC cells. We demonstrate for the first time that high stromal expression of SLC7A11 in human PDAC tissues predicts poorer overall survival. SLC7A11 inhibition in human patient-derived CAFs reduces their proliferation, anti-oxidant capacity and ability to support PDAC cell proliferation in 3D co-cultures. We also demonstrate the therapeutic potential of inhibiting SLC7A11 expression using a potent and selective gene silencing nanomedicine to decrease orthotopic PDAC tumour growth, stellate cell activation and fibrosis.

## RESULTS

### High stromal SLC7A11 expression in human PDAC specimens predicted poor overall survival

Results showed that SLC7A11 and its partner SLC3A2 mRNA levels are upregulated in PDAC patient-derived CAFs compared to patient-derived non-cancerous pancreatic fibroblasts (**Figure 1A**). Similar results were obtained when we analysed Ohlund et al data (25), which showed SLC7A11 mRNA expression was increased in iCAF (2.7-fold) and myCAF (1.6-fold) sub-populations compared to quiescent pancreatic stellate cells (pancreatic stellate cells differentiate into CAFs; **Supplementary Table 1**). We confirmed that all human PDAC CAFs expressed SLC7A11 protein (**Figure 1B**). SLC7A11 protein levels in human CAFs (5/6 CAFs tested) were comparable to the PDAC cells with the highest SLC7A11 expression (HPAFII and ASPC1) (**Figure 1B**). SLC7A11 protein levels were higher in PDAC cells derived from metastatic sites (HPAFII/AsPC1) relative to those from primary pancreatic tumours (MiaPaCa-2/Panc-1) (**Figure 1B**). Co-immunofluorescence staining for SLC7A11 and αSMA (CAF marker) in human PDAC tumours demonstrated abundant SLC7A11 protein in αSMA positive CAFs and tumour elements (**Figure 1C**). We next scored SLC7A11 protein expression, as determined by immunohistochemistry using a validated antibody (refer to methods and **Figure S1**), in tumour and stromal compartments of human PDAC tissue microarrays (**Figure 1D;** independent scoring scales were used for tumour and stromal compartments) and correlated with overall patient survival (**Figure 1E-G**). 58% of patients were Tumour^high^ (**Figure 1E**) and 47% were Stroma^high^ (**Figure 1F**). SLC7A11 expression in the PDAC tumour compartment alone did not predict patient survival (**Figure 1E**). Similar results were obtained when we analysed the ICGC publicly available mRNA data (26) (**Figure S2**). Importantly, in a multivariate logistic regression, no baseline variables were associated with stromal SLC7A11 expression and high SLC7A11 in the stroma was independently prognostic of poorer overall survival (**Figure 1F**, p=0.041, Hazard ratio=1.45; see **Supplementary Table 2** contains multivariate parameters), when adjusted for vascular invasion. In addition, we identified a sub-group of patients (Tumour^low^Stroma^high^) that had significantly poorer overall survival compared to all other score combinations (**Figure 1G-H**).

**Figure 1:**
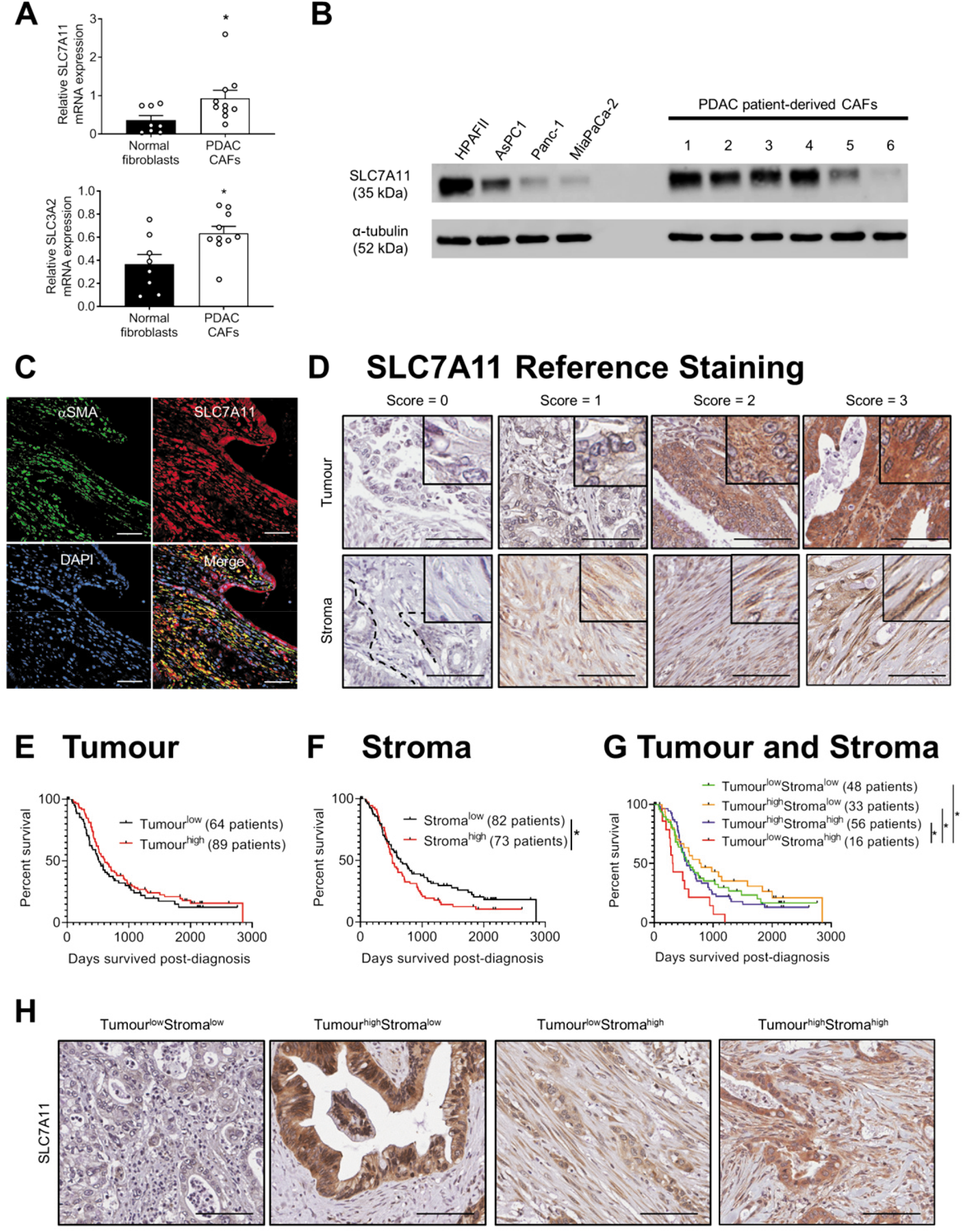
SLC7A11 is upregulated in human CAFs and can predict poorer overall survival in human PDAC patients. A) Quantitative real-time PCR analysis of SLC7A11 and SLC3A2 expression in total RNA extracts from normal pancreatic fibroblasts (isolated from n=8 patients with benign pancreatic conditions) and CAFs (isolated from n=10 PDAC patients). Bars show mean+s.e.m., circles indicate independent replicates (*p≤0.05, student t-test). B) Western blot of SLC7A11 in total protein extracts from PDAC cells and human CAFs (Cell lines 1-6). α-tubulin was used as a loading control. C) Immunoflourescence for DAPI, αSMA (CAF marker) and SLC7A11 in a human PDAC tissue specimen obtained through the Australian Pancreatic Cancer Genome Initiative (APGI). (D-G) Human PDA tissue microarrays obtained through the APGI (International Cancer Genome Consortium cohort) were stained for SLC7A11 by immunohistochemistry. D) Samples selected as references for scoring (0,1,2,3) for tumour and stromal compartments are shown (insets show magnified view of cells). Scores of 0-1 were classified as low SLC7A11 expression (“Tumour^low^” and “Stroma^low^”), scores of 2-3 were classified as high SLC7A11 expression (“Tumour^high^” and “Stroma^high^”). E-G) Kaplan-Meier survival curves showing the correlation between SLC7A11 expression in tumour cells (E), stroma (F), or a combination of both (G) with overall patient survival (days survived post-diagnosis). Patients that were deceased due to other causes or that were still alive were censored (shown as black ticks on each line graph). Total patient numbers for each group are indicated in the graph keys. Asterisks indicate significance based on (F) multivariate analysis (G) univariate Log-Rank test (*p≤0.05). H) Representative photos of Tumour^low^Stroma^low^, Tumour^high^Stroma^low^, Tumour^low^Stroma^high^, and Tumour^high^Stroma^high^ groups. Scale bars in all photos = 100μm.

### Inhibition of SLC7A11 in CAFs reduced cell proliferation and metabolically rewires CAFs

To assess SLC7A11 function in CAFs, we used siRNA and two pharmacological inhibitors [Sulfasalazine (SSZ), Erastin]. SLC7A11-siRNA potently reduced mRNA (**Figure S1C**) and protein levels (**Figure S1D**) compared to control siRNA. Inhibition of SLC7A11 by siRNA, SSZ or erastin significantly decreased CAF proliferation and viability (**Figure 2A-C**). Importantly, SLC7A11 knockdown in CAFs inhibited proliferation in both SLC7A11^low^ and SLC7A11^high^ cells (**Figure S3A**)], indicating that SLC7A11 is functionally essential in CAFs regardless of expression level. 2-mercaptoethanol (2-ME), which facilitates bypass of xCT for intracellular cysteine (27), rescued CAF growth in the presence of SSZ (**Figure 2B)**. This indicated that SSZ-induced growth arrest of CAFs was due to cystine starvation. To prove this, we assessed cystine uptake following SLC7A11 inhibition. Indeed, treatment of CAFs with either SLC7A11-siRNA or SSZ significantly reduced cystine uptake (**Figure 2D-E**) and intracellular glutathione (**Figure 2F-G**) relative to controls. The reduction in glutathione was rescued by addition of N-acetyl-cysteine (NAC; **Figure 2G**), which provides an SLC7A11-independent source of cysteine.

**Figure 2:**
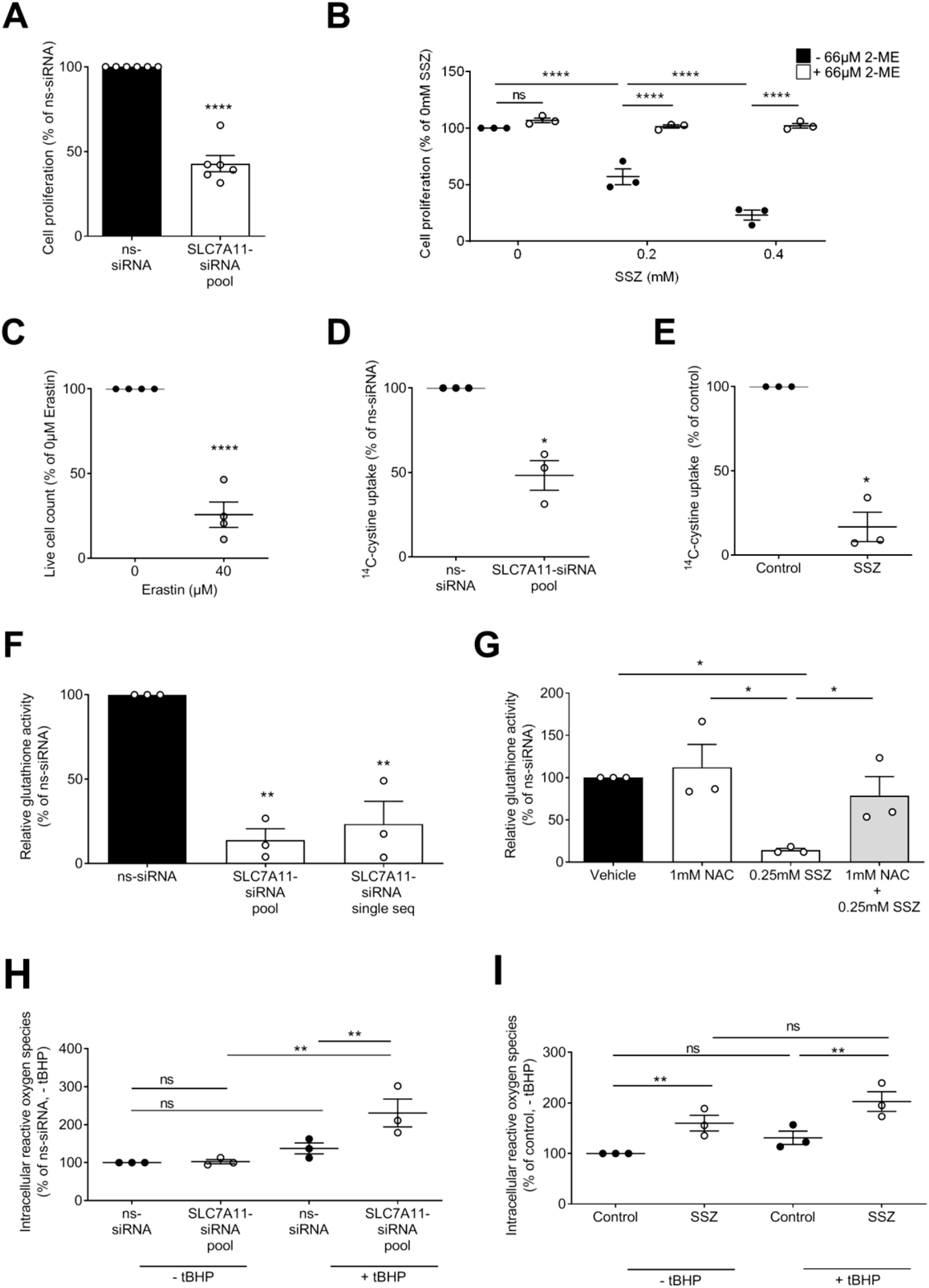
SLC7A11 inhibition in CAFs reduces proliferation and antioxidant capacity by inhibiting cystine uptake and glutathione production. A) Cell proliferation (based on cell counting kit 8 absorbance) of CAFs 72h post-transfection with non-silencing siRNA (ns-siRNA) or SLC7A11-siRNA pool (pool of 4 siRNA sequences). Asterisks indicate significance (****p≤0.0001; n=6, student t-test). B) Cell proliferation (cell counting kit 8 absorbance) of CAFs treated with sulfasalazine (SSZ) ± 66μM 2-mercaptoethanol (2-ME), as a % of controls. Circles indicate replicates, asterisks indicate significance (***p≤0.001, ****p≤0.0001; One-way ANOVA). C) Live cell counts of CAFs (as a fraction of controls) treated with erastin for 48h. Asterisks indicate significance (****p≤0.0001, n=4; student t-test). D-E) Radiolabelled cystine uptake as a fraction of ns-siRNA (72h post-transfection) or untreated control cells (48h post-treatment with SSZ). Asterisks indicate significance (*p≤0.05; student t-test). F-G) Intracellular glutathione levels as assessed by colorimetric assay, and as a fraction of ns-siRNA (72h post-transfection) or untreated control cells [16h post-treatment with SSZ±N-acetyl-cysteine (NAC)]. Asterisks indicate significance (*p≤0.05, **p≤0.01, n=6; One-way ANOVA). H-I) Intracellular oxidative stress in the presence or absence of tert-butyl hydroperoxide (tBHP; oxidative stress), as measured by CellROX staining and flow cytometry (as a fraction of ns-siRNA + 0 μM tBHP). Asterisks indicate significance (ns=not significant, **p≤0.01, n=3; One-way ANOVA). Circles in all graphs indicate replicates, lines and bars in all graphs represent mean±s.e.m.

We also validated previous findings that both SSZ and erastin significantly decreased MiaPaCa-2 PDAC cell proliferation (**Figure S3B-C**) and that SSZ inhibition can be rescued by 2-ME (17). In contrast to results in PDAC cells and CAFs, SLC7A11 knockdown had minimal effect on the viability (<20% reduction in viability) of non-tumour human pancreatic ductal epithelial cells (**Figure S3D-E**).

We next assessed intracellular reactive oxygen species (ROS; oxidative stress) and found that SLC7A11 knockdown in CAFs had no effect on intracellular ROS in the absence of stress, but significantly increased intracellular ROS in the presence of tBHP (**Figure 2H;** tBHP increased mitochondrial ROS, **Figure S4A**), suggesting decreased anti-oxidant capacity. In contrast, SSZ treatment alone increased intracellular ROS in CAFs, to levels where tBHP treatment had no significant additive effect on intracellular oxidative stress (**Figure 2I**). Note that the lack of an increase in intracellular ROS with tBHP alone is due to the short incubation time (1 hour) utilised for this assay. SLC7A11 knockdown in CAFs had no effect on glutamate secretion (**Figure S4B**).

### Inhibition of SLC7A11 increased sensitivity to oxidant stress and ferroptosis

Next, we wanted to examine whether the increase in intracellular ROS in CAFs makes them vulnerable to external oxidant stress (a common feature of the PDAC tumour microenvironment). We confirmed that SLC7A11 knockdown was maintained in the presence of oxidative stress (**Figure S4C**) and observed that oxidative stress increased SLC7A11 protein expression in cells treated with control siRNA (**Figure S4C**). Knockdown of SLC7A11 using siRNAs in CAFs sensitised them to external oxidant stress (tert-butyl hydroperoxide, tBHP) by decreasing viability (**Figure 3A**) and increasing apoptosis (**Figure 3B, Figure S4D**). Previous studies inhibiting SLC7A11 in PDAC cells have identified a key anti-proliferative mechanism to be induction of oxidative stress-induced cell death referred to as ferroptosis (19, 21, 28). We observed that inhibition of SLC7A11 with erastin reduced glutathione peroxidase activity indicative of ferroptosis in CAFs (**Figure 3C**). Importantly, the ferroptosis inhibitor ferrostatin rescued CAFs from the anti-proliferative effects of erastin further confirming ferroptosis (**Figure 3D**). Stable knockdown of SLC7A11 (**Figure S1F-G**) in CAFs using shRNA also significantly decreased cell viability in the presence of oxidant stress (**Figure 3E**) and increased ferroptosis (i.e. decreased glutathione peroxidase activity, **Figure 3F**).

**Figure 3:**
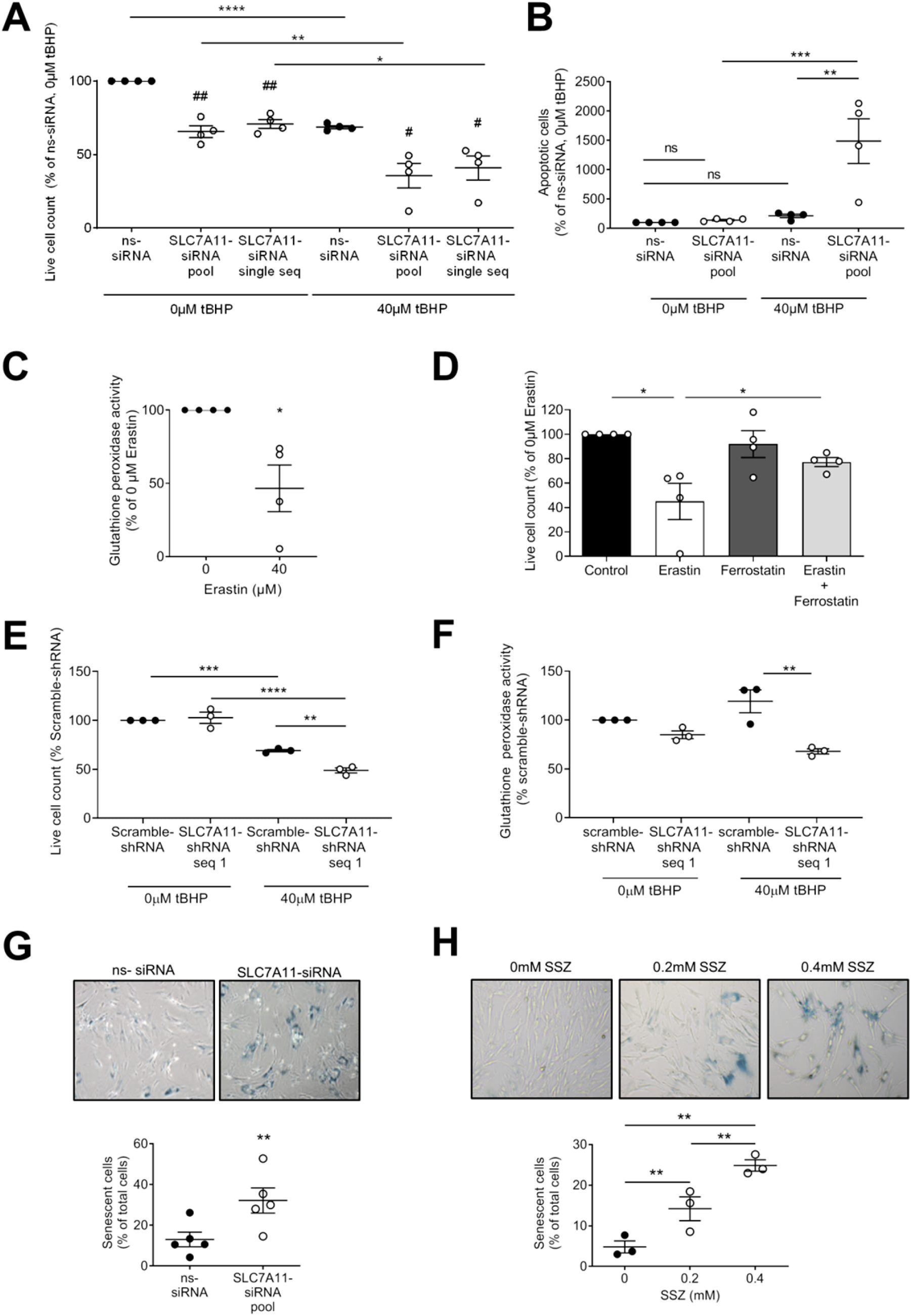
SLC7A11 inhibition in CAFs induces senescence and increases sensitivity to oxidative stress-induced cell death. A) Live cell counts of CAFs 72h post-transfection with non-silencing-siRNA (ns-siRNA), SLC7A11-siRNA pool or SLC7A11-siRNA single sequence (SLC7A11-siRNA single seq) and 24h post-treatment with tert-butyl hydroperoxide (tBHP). Circles indicate replicates, asterisks and hashes indicate significance (*p≤0.05, ** p≤0.01, **** p≤0.0001; # p≤0.05, ## p≤0.01, relative to ns-siRNA of the same tBHP concentration; n=4; One-way ANOVA). B) Frequency of AnnexinV+DAPI positive (apoptotic) cells, as a fraction of ns-siRNA+0μM tBHP controls, 72h post-transfection and 24h post-tBHP treatment. Circles indicate replicates, asterisks indicate significance (ns=not significant, **p≤0.01, ***p≤0.001; One-way ANOVA). C) Glutathione peroxidase activity of CAFs treated with erastin (9h) as a % of controls. Circles indicate replicates, asterisks indicate significance (*p≤0.05; n=4; student t-test). D) Live cell counts of CAFs 24h post-treatment with 40 μM erastin ± 2μM ferrostatin. Circles indicate replicates, asterisks indicate significance (*p≤0.05, n=4; One-way ANOVA). E) Live cell counts (trypan blue exclusion) of CAFs stably expressing scramble-shRNA or SLC7A11-shRNA seq 1, 72h post-seeding and 24h post-treatment with tBHP. Circles indicate replicate experiments. Asterisks represent significance (**p≤0.01, ***p≤0.001, ****p≤0.0001; n=3; One-way ANOVA). F) As per C, except CAFs stably expressed scramble-shRNA or SLC7A11-shRNA sequence 1 and were treated with 40uM tBHP for 9h, instead of erastin (**p≤0.01; n=3; One-way ANOVA). G-H) β-galactosidase positive cells (senescent cells) as a fraction of total cells (mean+s.e.m.): (G) 72h after transfection with control siRNA (ns-siRNA) or SLC7A11-siRNA or (H) 48h post-treatment with SSZ. Circles indicate replicates, asterisks indicate significance (**p≤0.01; G: n=4, student t-test; H: n=3, One-way ANOVA). Bars and lines in all graphs are mean±s.e.m. Replicate numbers in all panels refer to experiments performed using independent CAF cells isolated from different PDAC patients.

### SLC7A11 inhibition increased senescence of CAFs (in the absence of stress)

Given SLC7A11 siRNA alone had no effect on apoptosis (**Figure 3B**), we explored other anti-proliferative mechanisms. We showed that SLC7A11 siRNA had no effect on autophagy (**Figure S4E**), but increased CAFs in S-phase of cell cycle (**Figure S4F**, suggesting hindered S phase progression. Furthermore, we showed that SLC7A11 knockdown in CAFs significantly increased senescence (**Figure 3G**). This effect was reproduced by treatment of CAFs with SSZ (**Figure 3H**). Together our results showed that SLC7A11 knockdown in CAFs induced senescence and in the presence of additional oxidative stress compromised CAF survival.

### SLC7A11 inhibition decreased CAF and PDAC co-culture spheroid growth *in vitro*

To determine whether SLC7A11 inhibition in CAFs had any effect on their ability to support PDAC cell growth we performed 3D co-culture assays [spheroid outgrowth (**Figure 4A**) and spheroid growth assays (**Figure 4C**)]. Knockdown of SLC7A11 in either CAFs, PDAC cells, or both cell types significantly reduced spheroid outgrowth (**Figure 4B**). Importantly, knockdown of SLC7A11 in CAFs alone or in both cell types was more effective at inhibiting spheroid outgrowth than SLC7A11 knockdown in PDAC cells alone (**Figure 4B**). Using a stable knockdown approach in a 3D matrigel-embedded spheroid assay, we observed similar results (**Figure 4D**). Except SLC7A11-shRNA in MiaPaCa-2 PDAC cells alone had no effect on spheroid growth. In contrast, SLC7A11-shRNA in CAFs alone or in both tumour cells and CAFs reduced spheroid growth rate by > 30% (**Figure 4D**).

**Figure 4:**
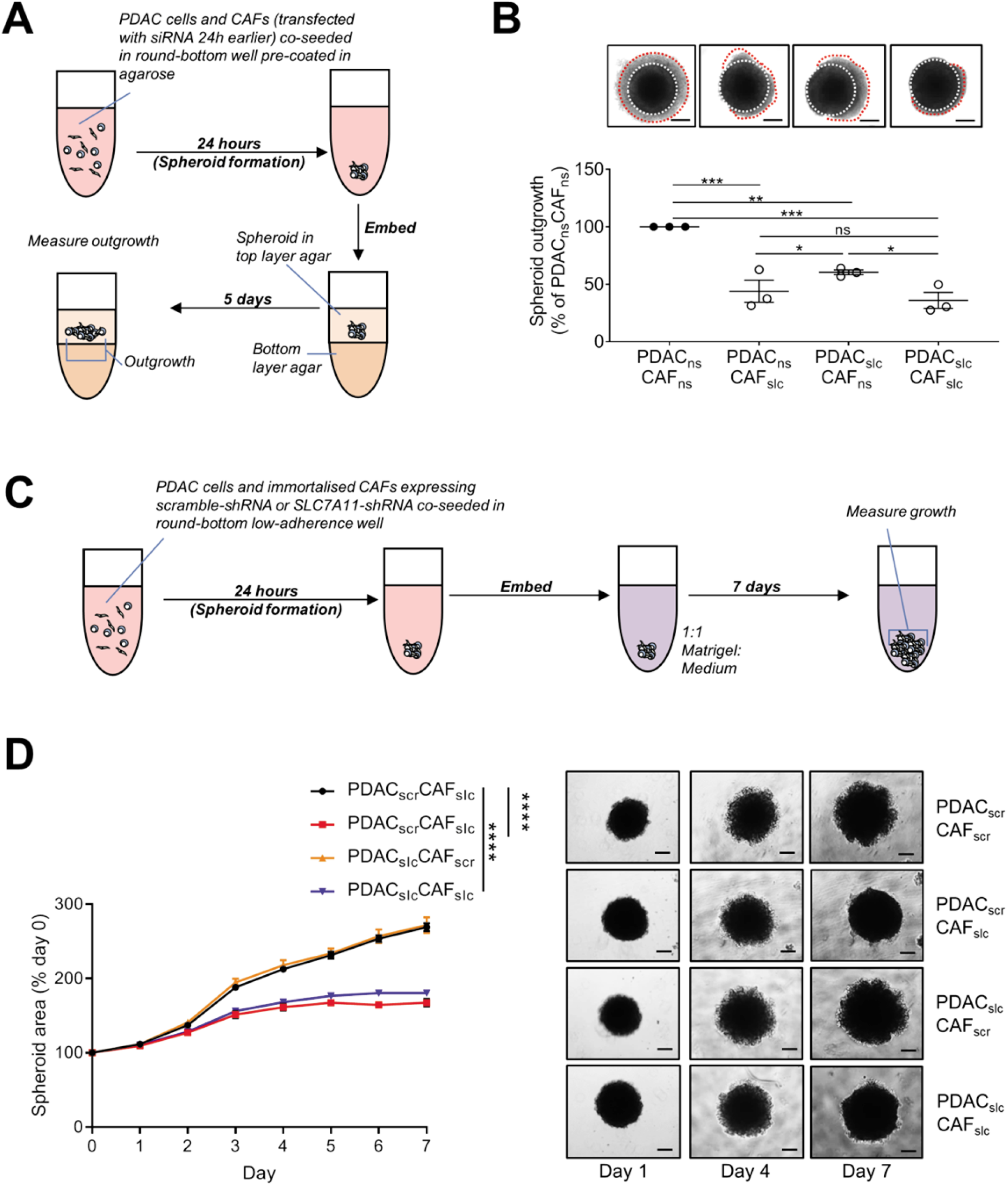
SLC7A11 knockdown in CAFs reduces pro-tumour cross-talk with PDAC cells in 3D co-culture spheroids. A-B) Schematic diagram of 3D co-culture spheroid outgrowth assay and quantification of 3D co-culture spheroid outgrowth post-transfection with control-siRNA (ns-siRNA) or SLC7A11-siRNA pool. Labels: **PDAC_ns_CAF_ns_** = non-silencing controls; **PDAC_ns_CAF_slc_** = SLC7A11 knockdown in CAFs only; **PDAC_slc_CAF_ns_** = SLC7A11 knockdown in PDAC cells only; **PDAC_slc_CAF_slc_** = SLC7A11 knockdown in both PDAC cells and CAFs. Representative photos are shown above each bar with the core circled in white dashed lines and outgrowth in red dashed lines (bars in photos = 300μm). Circles indicate replicates, lines indicate mean±s.e.m., asterisks indicate significance (ns=not significant, *p≤0.05, **p≤0.01, ***p≤0.001; One-way ANOVA). C) Schematic diagram of 3D co-culture growth assay using stable shRNA cell lines and D) representative photos (bars in photos = 200μm) and quantification of 3D co-culture spheroid growth. Labels: **PDAC_scr_CAF_scr_** = scramble-shRNA controls; **PDAC_scr_CAF_slc_** = SLC7A11-shRNA seq 1 in CAFs only; **PDAC_slc_CAF_scr_** = SLC7A11-shRNA seq 1 in PDAC cells only; **PDAC_slc_CAF_slc_** = SLC7A11-shRNA seq 1 in both PDAC cells and CAFs. Circles indicate replicates, lines indicate mean±s.e.m., asterisks indicate significance (ns=not significant, *p≤0.05, **p≤0.01, ****p≤0.0001, n=4-6; One-way ANOVA). Replicate numbers in panel B refer to independent experiments performed using MiaPaCa-2 combined with CAF cells isolated from different PDAC patients. Replicate numbers in panels D refer to replicate spheroids performed using MiaPaCa-2 PDAC cells combined with an immortalised CAF line.

### SLC7A11 inhibition decreased collagen remodelling *in vitro*

To examine the impact of SLC7A11 knockdown in CAFs on ECM remodelling, we used a matrix contractility assay (higher contraction = greater remodelling; **Figure 5A**). SLC7A11 knockdown in CAFs significantly reduced contraction of collagen plugs over 6 days (**Figure 5A**). Brightfield analysis of picrosirius red-stained collagen plugs at the end of the assay demonstrated that collagen plugs remodelled by CAFs transfected with SLC7A11-siRNA had decreased collagen relative to controls (**Figure 5B**). Polarised light analysis (measures density of collagen fibrils) showed that plugs remodelled by CAFs transfected with SLC7A11-siRNA had less overall birefringent fibrils (**Figure 5C**), decreased high and medium density fibrils, and significantly increased low density fibrils, relative to controls (**Figure 5D**). This was also confirmed by Second Harmonics Generation (SHG) analysis of fibrillar collagen (**Figure 5E**). Fibril organisation was also assessed by Grey-Level Co-Occurrence Matrix (GLCM) analysis of SHG images but showed no significant difference between ns-siRNA and SLC7A11-siRNA groups (**Figure 5E**).

**Figure 5:**
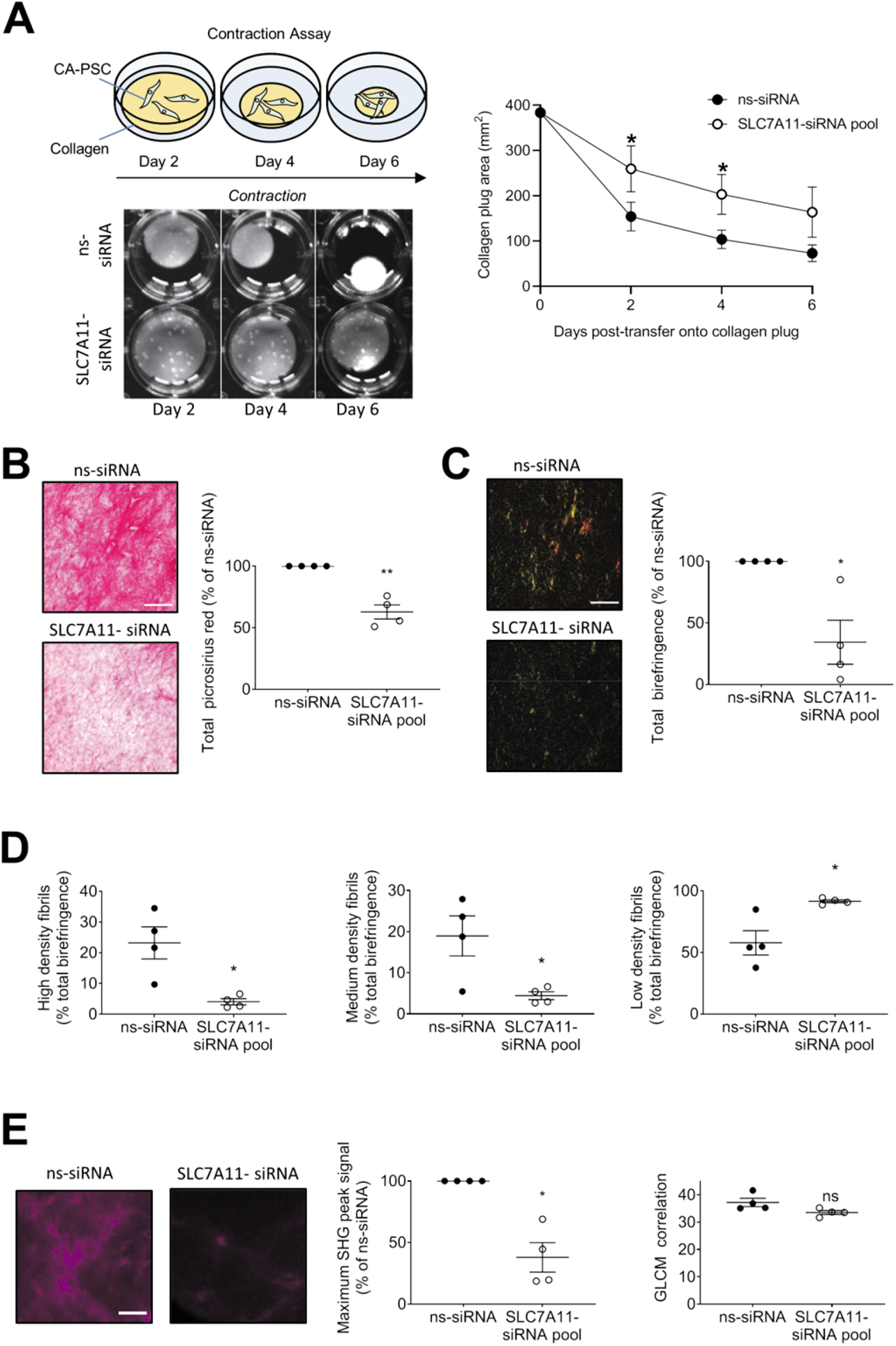
SLC7A11 knockdown in CAFs hinders collagen remodelling *in vitro*. A) Schematic diagram of assay and representative photos of collagen plugs contracted by CAFs transfected with control-siRNA (ns-siRNA) or SLC7A11-siRNA pool over 6 days are shown. The line graph shows the average area of contracted plugs at the indicated time points (mean±s.e.m.; *p≤0.05, n=4; One-way ANOVA). B-E) Analysis of collagen content in collagen plugs contracted by CAFs transfected with ns-siRNA or SLC7A11-siRNA at assay endpoint. B) Average picrosirius red signal. Circles indicate replicates, lines indicate mean±s.e.m., asterisks indicate significance (**p≤0.01, n=4; student t-test). Representative images of picrosirius red staining are shown. C) Average total birefringence. Circles indicate replicates, lines indicate mean±s.e.m., asterisks indicate significance (*p≤0.05, n=4; student t-test). Representative birefringence images are shown. D) Average % of total birefringence that was high (red-orange), medium (yellow) and low (green). Circles indicate replicates, lines indicate mean±s.e.m., asterisks indicate significance (*p≤0.05, n=4; student t-test). E) Left graph shows the average maximum second harmonics generation (SHG) signal detected by two-photon confocal microscopy of collagen plugs. Representative SHG images are shown. Right graph shows the average correlation based on GLCM analysis of SHG maximum intensity projections. Circles indicate biological replicates, lines indicate mean±s.e.m., asterisks indicate significance (ns=not significant, *p≤0.05, n=4; student t-test). Replicate numbers in all panels refer to independent experiments performed using independent CAF cells isolated from different PDAC patients.

### Genetic ablation of SLC7A11 in PDAC cells only had no effect on tumour growth in genetically engineered mouse models

Results above suggested that the presence of CAFs would likely influence the effect of an SLC7A11 inhibition approach *in vivo*. We assessed the impact of genetic ablation of SLC7A11 (**Figure S5**) driven by a pancreas-specific promoter (*Slc7a11^fl/fl^*, does not affect CAFs), in transgenic mouse models of PDAC (KC and KPC mice (29)). We observed no significant difference in PDAC precursor lesion formation between control and *Slc7a11^fl/fl^* KC mice (**Figure 6A**). In addition, in KPC mice we did not observe a significant difference in survival (**Figure 6B**) or intratumoural αSMA-positive CAFs (**Figure 6C**), but we did note a significant decrease in intratumoural collagen in KPC *Slc7a11^fl/fl^* mice, relative to controls (**Figure 6D**). We subsequently tested the effect of SLC7A11-siRNA on isolated KPC PDAC cells and KPC CAFs *in vitro*. Higher basal expression of SLC7A11 was observed in KPC PDAC cells relative to CAFs (**Figure S6A**) and siRNA potently knocked down SLC7A11 in both cell types (**Figure S6B-C**). Consistent with *in vivo* results, inhibition of SLC7A11 using an siRNA transient orthogonal approach had no effect on KPC PDAC cell proliferation (**Figure 6E**). However, SLC7A11 siRNA did significantly reduce proliferation of KPC CAFs (**Figure 6F**).

**Figure 6:**
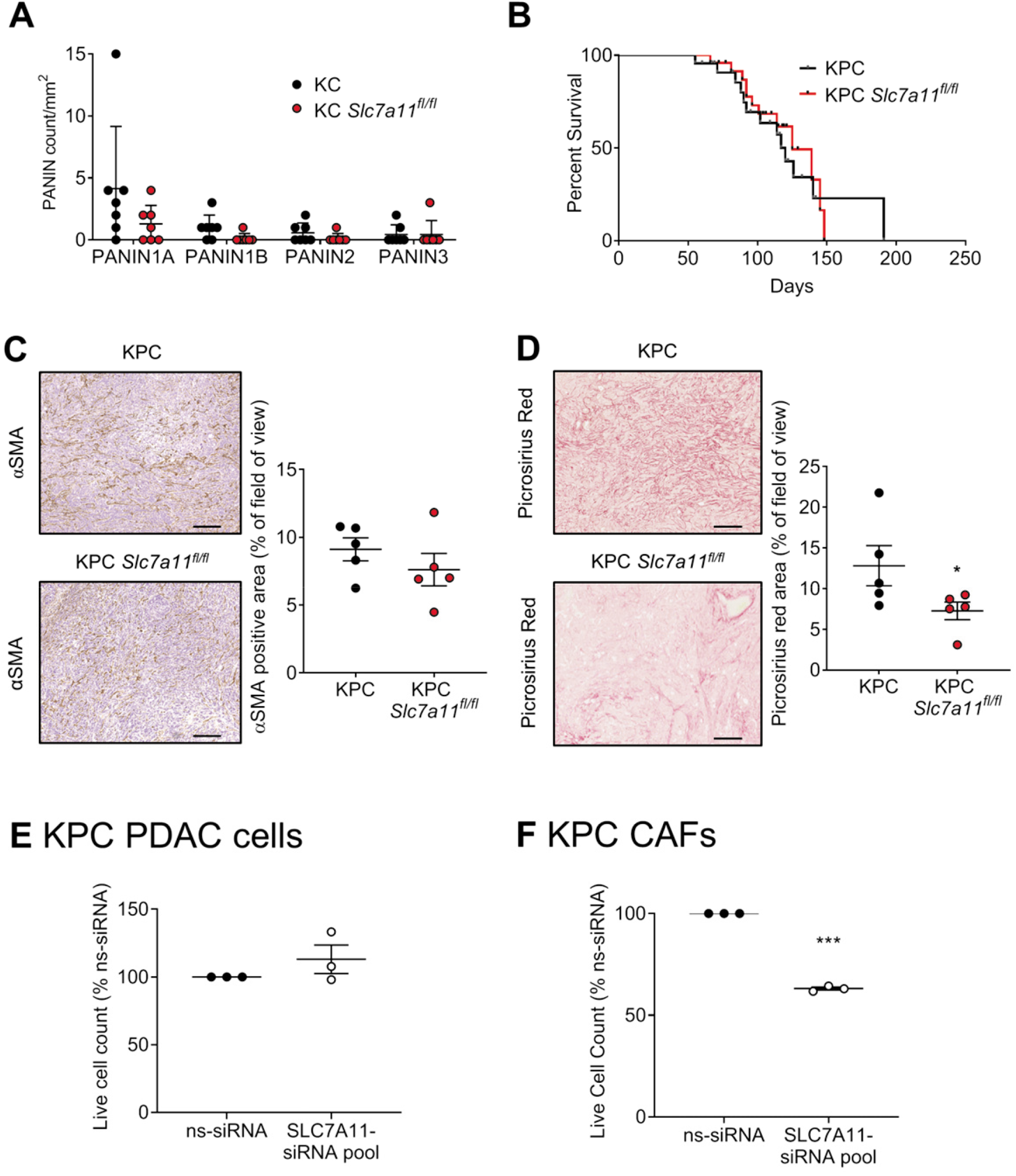
Genetic ablation of SLC7A11 in PDAC cells had no effect on orthotopic pancreatic tumour growth *in vivo*. A) Quantification of Pancreatic Intraepithelial Neoplasia (PanINs) 1A-3 from KC mice (n=7) and KC mice with SLC7A11 conditional KO under *Pdx1*-promoter (KC *Slc7a11^fl/fl^*; n=7) at 70 days of age (mean±s.e.m.). B) Kaplan-Meier analysis showing survival percentage of KPC (n=26) and KPC *Slc7a11^fl/fl^* mice (n=24) mice. C) Representative photos of KPC and KPC *Slc7a11^fl/fl^* tumour sections probed for αSMA (brown). The quantification of αSMA staining is shown in the graph (mean±s.e.m.), based on ImageJ analysis of representative regions from each tumour section (n=5 mice per group). Scale bars = 400μm. D) Representative photos of KPC and KPC *Slc7a11^fl/fl^* tumour sections. The quantification of picrosirius red staining is shown in the bar graph (mean±s.e.m.), based on ImageJ analysis of representative regions from each tumour section. Scale bars = 400μm. Asterisks indicate significance (*p≤0.05; n=5 mice per group; student t-test). E-F) Live cell counts (mean±s.e.m.) of (E) KPC PDAC cells and (F) KPC CAFs 72h post-transfection with control-siRNA (ns-siRNA) or mouse SLC7A11-siRNA pool (SLC7A11 pool). Asterisks indicate significance (*p≤0.05, **p≤0.01; n=3; one-way ANOVA).

### Gene silencing nanoparticles targeting SLC7A11 decreased orthotopic tumour growth, CAF activity and fibrosis

To overcome the physical barrier of fibrosis and deliver therapeutics to PDAC mouse tumours we developed di-block polymeric nanoparticles (Star 3), which can self-assemble therapeutic siRNA to form a nanocomplex that is stable in circulation and can extravasate from tumour vessels (30). Star 3-SLC7A11-siRNA decreased SLC7A11 protein levels in orthotopic pancreatic tumours (Figure 7A). The therapeutic efficacy of Star 3-SLC7A11-siRNA against orthotopic pancreatic tumours was then assessed using a therapeutic regimen, with or without co-administration of Abraxane^®^ (human albumin-bound paclitaxel; currently used in combination with gemcitabine to treat PDAC in the clinic; Figure 7B). Whilst Abraxane^®^ treatment had no effect on tumour growth, SLC7A11 inhibition alone, or in combination, significantly decreased tumour growth (Figure 7C), and reduced the incidence of metastases (Table 1), but had no effect on the number of metastases per mouse (Figure 7D). Furthermore, Star 3-SLC7A11-siRNA significantly decreased the frequency of intratumoural αSMA positive cells (Figure 8A) and picrosirius red staining (fibrosis), relative to controls (Figure 8B), though fibril density and organisation were not significantly affected (Figure 8C; Figure S7A-B). Our results suggested that SLC7A11 knockdown reduced total intratumoural collagen rather than the quality of remaining collagen. This resulted in an increase in the fraction of open CD31-positive blood vessels, relative to controls (Figure 8D), suggesting normalisation of intratumoural vasculature.

**Table 1:** Metastases incidence in orthotopic PDAC model treated with Star 3+SLC7A11-siRNA.

| **Incidence of metastases** |  |  |  |
| --- | --- | --- | --- |
| **(% of mice with metastases)** |  |  |  |
| **Control-siRNA**<br>**+ Albumin** | **Control-siRNA +**<br>**Abraxane** | **SLC7A11-siRNA**<br>**+ Albumin** | **SLC7A11-siRNA**<br>**+ Abraxane** |
| **50** | **37.5** | **28.6** | **37.5** |

**Figure 7:**
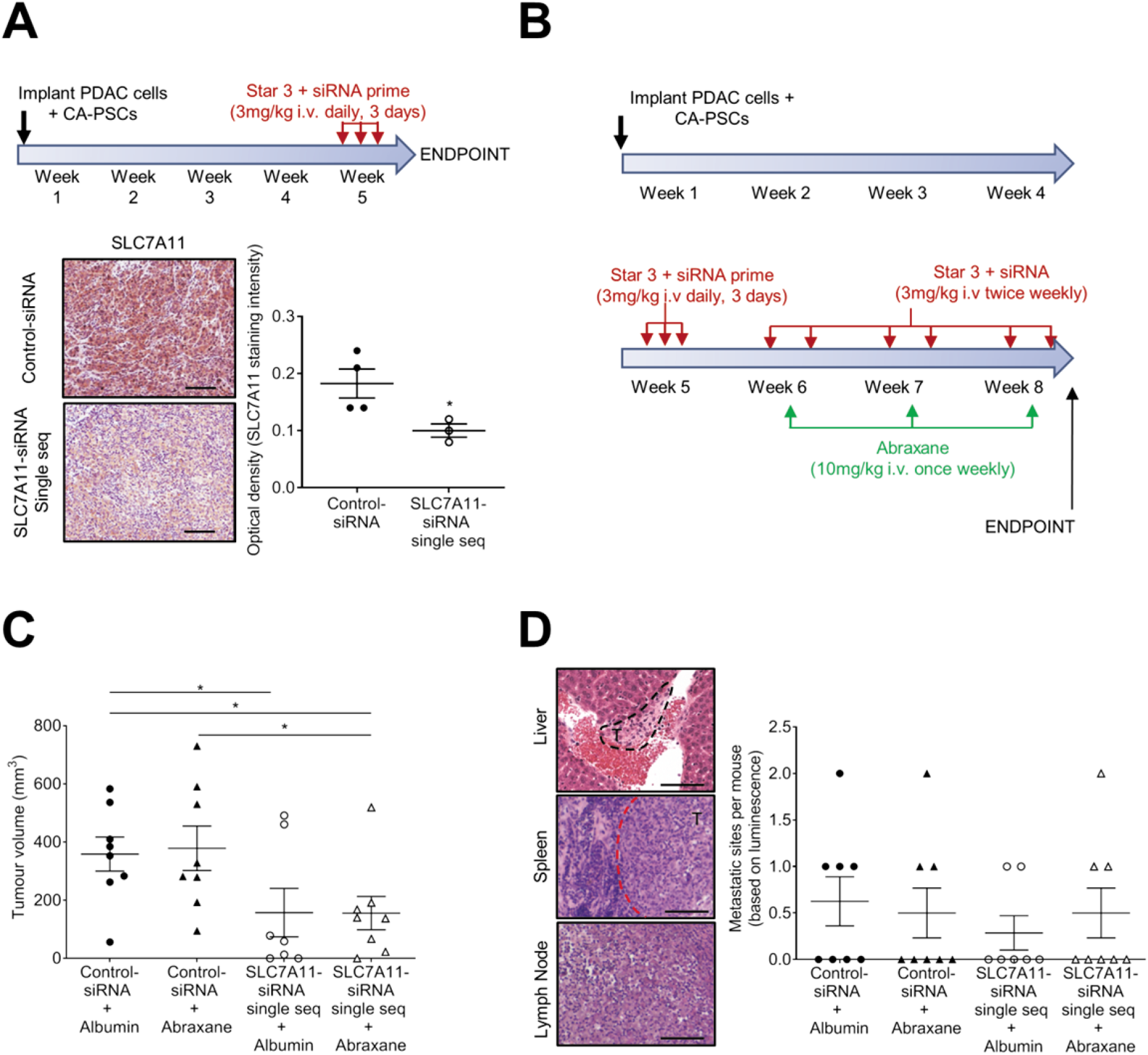
Star 3+SLC7A11-siRNA treatment reduces orthotopic pancreatic tumour growth and metastasis. All orthotopic tumours were co-injections of PDAC cells and CAFs. A) Orthotopic pancreatic tumours were treated with STAR nanoparticles + control-siRNA or SLC7A11 siRNA single sequence (SLC7A11 single seq) in the regimen shown. Representative photos of immunohistochemistry for SLC7A11 in tumour tissue at the model endpoint are shown. Graph shows optical density (staining intensity) calculated from average pixel intensity measurements from 3 representative images per tumour, using ImageJ. Circles indicate individual mice, lines indicate mean±s.e.m., asterisks indicate significance (*p≤0.05; One-way ANOVA). B) Treatment regimen for therapeutic model analysed in panels (C-D). Circles and triangles in all dot plots in panels C-D represent individual mice. C) Tumour volume at therapeutic model endpoint, as assessed by calliper measurement *ex vivo* (mean±s.e.m.). Asterisks indicate significance (*p≤0.05; One-way ANOVA). D) Representative photos of metastases confirmed by H&E staining following detection at model endpoint by *ex vivo* luminescence imaging of organs. Graph shows metastatic sites per mouse (mean±s.e.m.) for each treatment group. Scale bars in all figures = 200μm.

**Figure 8:**
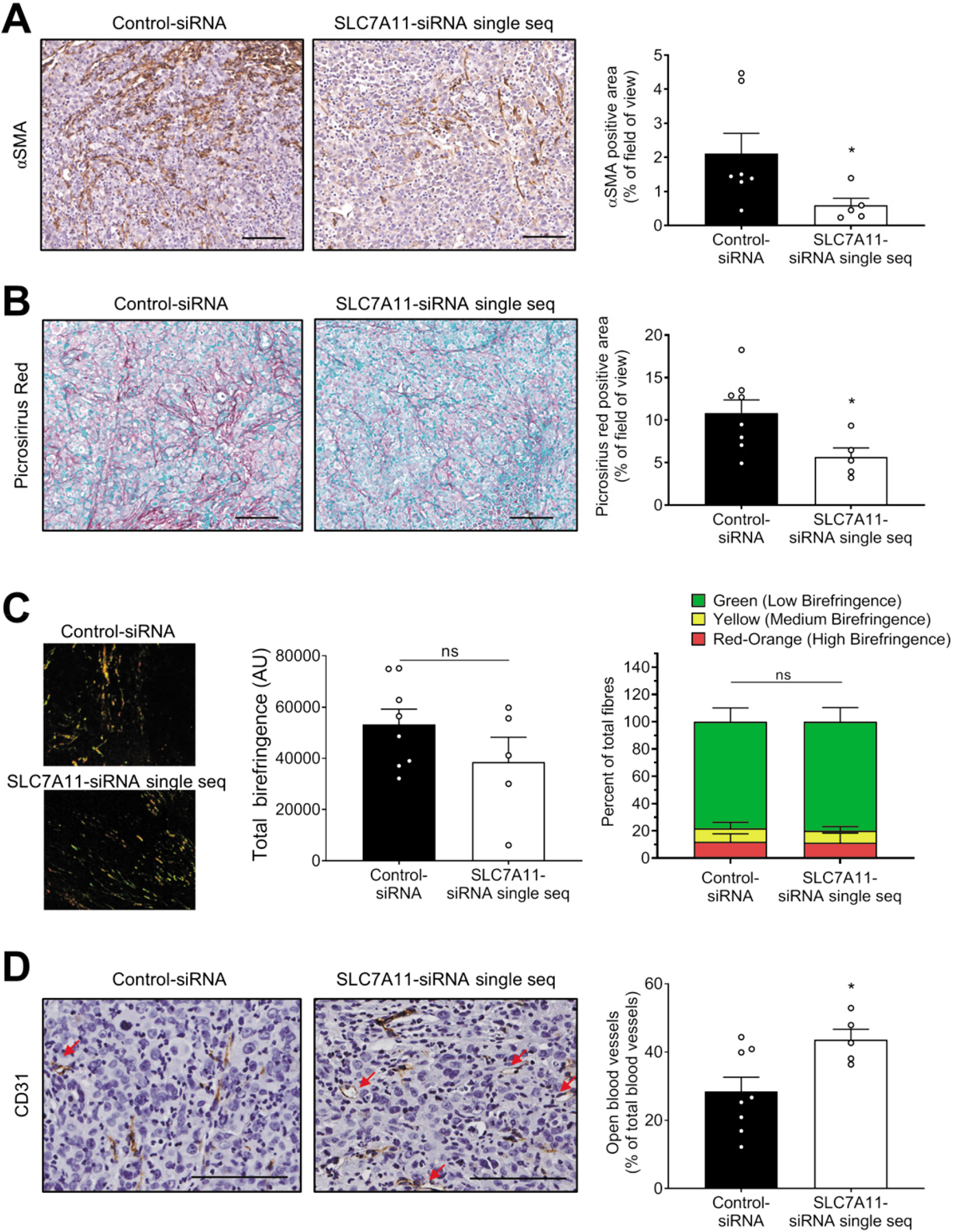
Star 3+SLC7A11-siRNA treatment of orthotopic pancreatic tumours reduces intratumoural CAF activation and fibrosis, and normalises tumour vasculature. A) Representative photos of tumour sections probed for αSMA (brown). The quantification of αSMA staining is shown in the graph (mean+s.e.m.), based on ImageJ analysis of representative regions from each tumour section. Asterisks indicate significance (*p≤0.05; Control-siRNA, n=7; SLC7A11-siRNA single seq, n=5; student t-test). B) Representative photos of picrosirius red and methyl green stained tumour sections. The quantification of picrosirius red staining is shown in the bar graph (mean+s.e.m.), based on ImageJ analysis of representative regions from each tumour section. Asterisks indicate significance (*p≤0.05; Control-siRNA, n=8; SLC7A11-siRNA single seq, n=5; student t-test). C) Polarised light analysis of representative regions from picrosirius red stained specimens. Representative photos are shown. Left bar graph shows total birefringence (mean+s.e.m.; Control-siRNA, n=8; SLC7A11-siRNA single seq, n=5). Right bar graph shows the average frequency (mean+s.e.m.; Control-siRNA, n=8; SLC7A11-siRNA single seq, n=5) of low, medium and high birefringence collagen fibrils (higher birefringence = denser fibril). ns = not significant (student t-test). D) Representative photos of CD31-stained tumour sections. Red arrows indicate open blood vessels. The bar graph shows the fraction of CD31-positive blood vessels that were open (mean+s.e.m.), based on ImageJ analysis of representative regions from each tumour section. Asterisks indicate significance (*p≤0.05; Control-siRNA, n=8; SLC7A11-siRNA single seq, n=5; student t-test). Fields of view used for analyses in all panels, provided an average area coverage of 13% of the total tumour section (excluding necrotic regions). All circles in graphs represent individual mice. All scale bars in photos = 100μm.

## DISCUSSION

PDAC urgently requires more effective treatments that target both tumour cells and the tumour/fibrosis-promoting stromal CAFs. In this study, we showed for the first time that high SLC7A11 expression in the stroma of human PDAC tumours predicts poorer patient survival. We also demonstrated that limiting cystine uptake in CAFs via inhibition or knockdown of the amino acid transporter SLC7A11, halted their proliferation and sensitised them to oxidative stress. In a 3D CAF/PDAC cell co-culture setting, this approach led to decreased spheroid growth, indicating that we had disrupted PDAC-CAF cross-talk. SLC7A11 knockdown in CAFs also decreased their ability to remodel 3D collagen *in vitro*, implying this approach had the potential to affect matrix remodelling in PDAC tumours. Finally, we showed that our siRNA-containing nanoparticles were able to decrease mouse tumour growth, incidence of metastases and fibrosis. A key finding of this study is the importance of SLC7A11 in regulating the growth and function of CAFs, and that inhibiting SCL7A11 in CAFs is essential to maximise the full therapeutic benefit of targeting SLC7A11 in PDAC.

High SLC7A11 expression has been reported to predict poorer survival in different tumour types (14, 31, 32). However, the role of SLC7A11 in PDAC is less clear. Maurer et al (33) and more recently Badgley et al (21) observed significantly higher levels of SLC7A11 mRNA in the PDAC epithelial compartment relative to stroma. Yang et al (34) recently showed that expression of the long non-coding RNA, SLC7A11-AS1 (antisense transcript of SLC7A11) was also increased in PDAC tumour tissue relative to normal pancreas, and higher levels of SLC7A11-AS1 among PDAC patients predicted poorer overall survival. SLC7A11-AS1 in PDAC cells was found to act as a scavenger for reactive oxygen species by preventing proteasome degradation of nuclear factor erythroid-2-related factor 2 (Nrf2), a key master regulator of redox homeostasis. The results add another layer of oxidative stress protection indirectly regulated by SLC7A11 expression. We analysed gene expression array data from the ICGC PACA-AU cohort and found that SLC7A11 mRNA expression did not correlate with overall survival. However, using an immunohistochemistry (IHC)-based approach in the ICGC cohort, we demonstrated that high SLC7A11 protein in the stroma, but not in the tumour compartment, was prognostic of poorer patient survival. The difference between expression array and IHC results might be explained by the lack of segregation of tumour and stroma in the ICGC gene expression array data set. This is a critical consideration, given the ICGC PDAC cohort has been identified as the most stroma-rich cohort in comparison to TCGA and UNC PDAC cohorts (33). In our IHC approach, tumour and stroma were scored on separate scales to account for differences in maximum expression between the compartments and to prevent the masking of stromal SLC7A11 expression by the higher average levels of expression in tumour elements. Interestingly, we identified a sub-population of patients with a combination of high stromal SLC7A11 expression and low tumour expression that had significantly poorer overall survival than all other PDAC patients. This might be indicative of metabolic cross-talk between PDAC cells and CAFs in a subset of patients. Studies have demonstrated that decreased SLC7A11 in tumour cells can confer resistance to glucose deprivation by increasing glutamate retention (35, 36), as glutamate can fuel the TCA cycle under low glucose conditions. Thus, a situation where PDAC tumour cells can increase glutamate retention, while potentially sourcing cysteine from nearby SLC7A11^high^ stromal cells, could be advantageous for their survival. Our results highlight that despite the higher average expression of SLC7A11 in the tumour compartment, expression in the PDAC stroma may be more functionally significant for disease progression. In addition, a retrospective analysis of expression array data from (25) showed that SLC7A11 expression was elevated in iCAFs and myCAFs relative to quiescent pancreatic fibroblasts, particularly in iCAFs. Future studies will investigate the potential role of SLC7A11 in the immune modulatory functions of iCAFs.

We used orthogonal approaches consisting of two pharmacological inhibitors [sulfasalazine (SSZ), erastin] and RNA interference (siRNA, shRNA) to show that inhibition of SLC7A11 in CAFs reduced CAF proliferation, and that this effect was abrogated by β-mercaptoethanol (2-ME). 2-ME bypasses the need for SLC7A11 by reducing cystine to cysteine, thus the need for SLC7A11 cystine shuttling becomes redundant (27). Ferrostatin (a ferroptosis inhibitor) was also able to prevent the decrease in CAF proliferation when exposed to erastin. These results are in support of a recent study by Badgley et al (21) which demonstrated that inhibition of SLC7A11 in PDAC cells induced ferroptosis. Interestingly both SLC7A11 knockdown and SSZ treatment in CAFs were found to induce senescence. This is the first time senescence has been reported as a response to SLC7A11 inhibition and may have been a survival response to amino acid deprivation and potentially hindered protein synthesis. Indeed, Daher et al (19) demonstrated that genetic ablation of SLC7A11 in PDAC cells induced an amino acid stress response.

We confirmed that both SLC7A11 knockdown and SSZ significantly hindered cystine uptake as well as the production of the antioxidant GSH. Given cystine is integrated into GSH as cysteine, and cysteine levels control GSH synthesis, these results implied that SLC7A11 inhibition decreased intracellular cysteine. A key point of difference between siRNA-based and SSZ-based approaches was that SLC7A11 knockdown alone did not significantly increase oxidative stress in CAFs, whereas SSZ did. A potential explanation for this difference is that SSZ can decrease levels of additional enzymes and signalling pathways involved in protection against oxidative stress (37-39), which might induce higher oxidative stress more rapidly than SLC7A11 gene silencing. Importantly, supplying cells with an SLC7A11-independent source of cyst(e)ine via N-acetyl-cysteine was able to rescue GSH levels in SSZ-treated CAFs. Our results highlighted the crucial dependence of CAFs on SLC7A11 for cystine uptake and GSH synthesis.

Our novel findings led us to investigate the potential impact of SLC7A11 inhibition on cross-talk between PDAC cells and CAFs. Importantly, in the spheroid growth assay, stable SLC7A11 knockdown in PDAC cells alone had no effect on spheroid growth. This lack of an effect may have been due to the reduced contribution of PDAC cells to the total spheroid volume in the spheroid growth assay (3:1 excess of CAFs), as opposed to the equal ratio of PDAC cells:CAFs used in the outgrowth assay. In addition, the presence of a basement matrix in the spheroid growth assay may have contributed to the phenotype, as CAF-mediated matrix remodelling would have assisted spheroid local invasion and growth. SLC7A11 knockdown in CAFs also significantly reduced their ability to remodel 3D collagen *in vitro*, suggesting that SLC7A11 inhibition in CAFs might remodel a key barrier to drug delivery *in vivo*. Our results reiterate the importance of targeting SLC7A11 in both CAFs and PDAC cells.

Consistent with this, conditional knockout of SLC7A11 in only the tumour compartment of KPC tumours did not affect mouse survival, but interestingly it did decrease intra-tumoural fibrosis, implying pro-fibrogenic cross-talk between tumour and stromal cells had been disrupted. Notably, these results were reproduced *in vitro*, whereby SLC7A11 knockdown in isolated CAFs from the KPC mouse tumours significantly reduced their proliferation but had no effect in KPC PDAC cells. Our results are in striking contrast to prior KPC mouse PDAC models in which SLC7A11 was inhibited via a modified form of erastin (20), systemic deletion of SLC7A11 or cysteinase enzyme treatment (21) and significantly reduced tumour growth. What is important to note is that these studies did not selectively target PDAC tumour cells and these mouse models are characterised as having a prominent fibrotic tumour stroma. Therefore, it is likely that SLC7A11 inhibition would have also affected CAFs pro-tumour/fibrotic activity which would have contributed to the reduced tumour growth.

Both prior studies also demonstrated that SLC7A11 inhibition (20) or systemic genetic deletion (21) in the KPC mouse model increased median survival, highlighting the efficacy of SLC7A11 inhibition as a standalone therapeutic approach. To complement this work, we opted for an orthotopic model of PDAC with a defined pre-mortality endpoint. Our approach had the advantage of utilising human-derived PDAC cells and CAFs and the defined endpoint allowed for a time-matched comparison of the effect of SLC7A11 knockdown on CAF activation, fibrosis and tumour size. Using this model, we tested the therapeutic efficacy of a polymeric nanoparticle (Star 3) that we specifically developed to package SLC7A11-siRNA and overcome physical barriers to drug delivery to penetrate fibrotic PDAC tumours in mice (30). Our nanoparticle preferentially accumulates in PDAC tumours, but is rapidly cleared from normal organs, minimising the chance for off-target toxicity (30). Star 3-SLC7A11-siRNA was able to decrease tumour growth by >60%. This was coupled with reduced incidence of metastases, decreased CAF activation and intratumoural collagen (fibrosis), as well as normalised tumour vasculature. It should be noted that our approach used human-specific SLC7A11-siRNA and would not have targeted mouse cells. Thus, the effects we observed may be an underestimate of the effect of SLC7A11 inhibition in PDAC tumours, as mouse CAFs can be co-recruited/activated in the model. Regardless, our results demonstrate the efficacy of Star 3-SLC7A11-siRNA against PDAC tumours and its ability to alleviate a physical barrier to drug delivery. While this did not translate into increased sensitisation of orthotopic PDAC tumours to Abraxane^®^, the dosing schedule selected was suboptimal to test whether SLC7A11 inhibition could sensitise to lower amounts of Abraxane^®^. Future studies will investigate the potential for sensitisation to higher doses of Abraxane as well as identify other potential drug/therapeutic combinations using drugs (i.e. gemcitabine, cisplatin, carboplatin) or irradiation which increase intracellular reactive oxygen species.

One limitation of our study was the lack of a proficient adaptive immune system in the host mice used. PDAC tumours have been demonstrated to be immune privileged, with immune infiltrate primarily consisting of immune suppressive M2 macrophages and regulatory T cells. A number of studies have now demonstrated that reprogramming of CAFs and the PDAC stroma can improve anti-tumour immune responses (40-42). Arensman et al (18) demonstrated that SLC7A11 was dispensable for T-cell proliferation and anti-immune response *in vivo*. Importantly, they showed that SLC7A11 knockout using CRISPR/cas9 gene editing in subcutaneous PDAC tumours sensitised them to anti-CTLA-4 immunotherapy (18), suggesting SLC7A11 inhibition might work in synergy with immunotherapies.

This study brings together over a decade of research into the therapeutic potential of SLC7A11 inhibition in PDAC. Taken together, our findings and those of previous studies have demonstrated that SLC7A11 inhibition in PDAC is a multi-pronged therapeutic approach that can reverse PDAC resistance by: (i) directly inhibiting PDAC cell (16-20) and CAF proliferation; (ii) increasing PDAC cell chemosensitivity (16, 17, 19); (iii) interfering with pro-tumour signalling and potentially nutrient exchange between PDAC cells and CAFs; (iv) alleviating a physical barrier to drug delivery (fibrosis); (v) enhancing anti-tumour immune responses (18).

## METHODS

### Quantitative real-time PCR (qPCR), Western blotting, siRNA transfections

Description is included in *Supplementary Materials and Methods*.

### Cell isolation and culture

Human PDAC cells (MiaPaCa-2, Panc-1, AsPC1 and HPAFII; American Tissue Culture Collection) were cultured as described (43-45). PDAC cell purity was confirmed by short tandem-repeat profiling (CellBank Australia). Normal Human Pancreatic Ductal Epithelial (HPDE) cells (a gift from Ming Tsao, Ontario Cancer Institute) were cultured in Keratinocyte-serum-free (KSF) medium containing 50 mg/ml bovine pituitary extract (BPE) and 5 ng/ml epidermal growth factor (EGF), as previously described (46). Quiescent human pancreatic fibroblasts, activated by culture on plastic, were isolated from patients with benign pancreatic conditions using a Nycodenz gradient centrifugation and cultured in IMDM containing 10% FBS and 4mM L-glutamine, as previously described (47). Human CAFs were isolated from PDAC tumour tissue by explant culture and cultured in IMDM containing 10% FBS and 4mM L-glutamine, as previously described (48, 49). The purity of CAFs was assessed by positive immunostaining for glial fibrillary acidic protein (GFAP) and alpha-smooth muscle actin (α-SMA) and negative immunostaining for cytokeratin, as described (47). All cells were maintained at 37°C in a humidified atmosphere containing 5% CO2 and were negative for mycoplasma. Cells were lifted by incubation in 0.05% trypsin (CAFs) or 0.25% trypsin (PDAC cells) and pelleted at 335 x g, 3 min at room temperature before resuspension.

### Immortalisation of human PDAC CAFs and establishment of human PDAC and CAF cell lines stably expressing shRNA

Human patient-derived PDAC CAFs at passage 9 (PSC line 1 in **Figure 1B**) were immortalised by lentiviral delivery of a human telomerase expression construct (GenTarget, Cat. LVP1131-RP). Cells were maintained in puromycin selection and red fluorescent protein positive cells sorted on a BD FACS Aria II cell sorter. MiaPaCa-2 cells and hTERT-immortalised CAFs were then transduced with lentiviral constructs expressing scramble shRNA, SLC7A11-shRNA sequence 1 (Origene, Cat. TL309282). Transduced cells were maintained in puromycin and GFP positive cells sorted on a BD FACS Melody cell sorter. SLC7A11 knockdown was confirmed by Western blot. All CAF shRNA cell lines were re-validated by positive immunostaining for GFAP and α-SMA, and negative immunostaining for cytokeratin.

### Isolation and culture of KPC transgenic mouse PDAC and CAF cells lines from PDAC tumours

KPC PDAC cells were supplied by co-authors Jen Morton and Paul Timpson and were cultured as previously described (50). KPC CAFs were isolated as previously described (48, 49) from KPC PDAC tumours and were validated by immunocytochemistry for GFAP and αSMA. KPC CAFs were cultured as per human PDAC CAF culture medium and conditions (48, 49).

### Immunofluorescence for SLC7A11 and αSMA co-localisation

Formalin-fixed, paraffin-embedded human PDAC tumour tissue was obtained through the Australian Pancreatic Cancer Genome Initiative. Antigen retrieval was performed as previously described (43-45). Tissue sections were then stained with the following antibodies: SLC7A11 (Cell Signalling Technologies, Cat. 12691; 1:25) and αSMA (Sigma-Aldrich, Cat. A5228; 1:1000) overnight at 4°C, followed by anti-rabbit-AF647 secondary antibody (Abcam Cat. ab150115) and anti-mouse-AF488 secondary antibody (Life Technologies, Cat. A11001; 1:1000) for 1h at room temperature. Tissues were then mounted using Prolong Gold Antifade mountant (ThermoFisher Scientific, Cat. P36931) and imaged on a Zeiss 900 confocal microscope.

### Immunohistochemistry comparison of SLC7A11 antibodies on human PDAC tumour tissue

Serial sections of formalin-fixed, paraffin-embedded human PDAC tumour tissue were obtained through the Australian Pancreatic Cancer Genome Initiative. Antigen retrieval was performed as previously described (43-45). Tissue sections were probed with the following antibodies: SLC7A11 (Cell Signalling Technologies, Cat. 12691, 1:25; Abcam, Cat. ab37185, 1:2000; Novus Biologicals, Cat. NB300-318, 1:2000), biotinylated anti-rabbit secondary antibody (Vector Laboratories, Cat. BA-1000; 1:50) and Vectastain^®^ ABC kit (Vector laboratories). 3,3’ diaminobenzidine was used as the substrate and hematoxylin as a counter-stain. Note that the cell signalling antibody used was validated based on requirements as detailed in (51). We showed similar staining patterns in PDAC tissue sections using 3 independent antibodies (**Figure S1A**). In addition, our positive control brain tissue (**Figure S1B**) had abundant SLC7A11 protein expression and our negative control skin tissue (**Figure S1B**) had no SLC7A11 expression, consistent with SLC7A11 expression levels defined in the human protein atlas. Specificity of the antibody was also confirmed by its ability to detect specific SLC7A11 gene silencing (siRNA and shRNA) in Western blot (**Figure S1D-G**).

### Correlation of SLC7A11 expression in human PDAC specimens with overall survival

#### Immunohistochemistry analysis

Formalin-fixed, paraffin-embedded human PDAC tissue microarrays (TMAs) were obtained through the Australian Pancreatic Cancer Genome Initiative (International Cancer Genome Consortium Cohort). Patient demographics are summarised in **Supplementary Table 3**. TMA rehydration and blocking for immunohistochemistry was performed as previously described (43-45). TMAs were probed with the following antibodies: SLC7A11 (Cell Signalling Technologies, Cat. 12691; 1:25), biotinylated anti-rabbit secondary antibody (Vector Laboratories, Cat. BA-1000; 1:50) and Vectastain^®^ ABC kit (Vector laboratories). 3,3’ diaminobenzidine was used as the substrate and hematoxylin as a counter-stain. Intensity of staining in tumour and stromal compartments was scored using a four-point scale (0-3) by two independent scorers, based on the intensity in ≥75% of each compartment (normal acinar and ductal cells not scored). Score scales for tumour and stroma compartments were independent of each other. A consensus score was obtained for each core. For each set of 3 cores per patient, the highest tumour and stroma scores were selected for correlation with patient parameters. Any non-PDAC tumours were excluded. Scores of 0-1 = SLC7A11^low^; Scores of 2-3 = SLC7A11^high^. Scores were then correlated with overall survival using a Kaplan Meier Survival Curve (see statistical analyses). Patients that were deceased due to other causes or still alive were censored. Note that 2 patients did not have a tumour compartment in all 3 cores and were excluded. ***RNA analysis:*** Normalised SLC7A11 expression values (expression array data) were from the PACA-AU cohort through the ICGC data portal (all expression data for the PACA-AU cohort publicly available at ICGC data portal: https://dcc.icgc.org/projects/PACA-AU). Non-PDAC patients were excluded (total PDAC patients = 242 patients). Normalised expression values were broken into tertiles (low = 0-1.98, medium = 1.98-2.52, high = 2.53-6.45) and correlated with overall survival. SLC7A11 mRNA expression was then correlated with overall survival using a Kaplan Meier Survival Curve (see statistical analyses). For comparison of SLC7A11 expression in mouse iCAFs, myCAFs and quiescent pancreatic fibroblasts (pancreatic stellate cells) normalised expression data from Ohlund et al (25). Details of experiments can be found in the original publication. Briefly, mouse pancreatic fibroblasts were isolated from wild-type C57Bl/6 mice by outgrowth and cultured as follows: (1) quiescent fibroblasts alone in Matrigel; (2) cultured in trans-well with tumour organoids = iCAFs (αSMA^low^IL-6^high^); (3) grown in monolayer = myCAFs (αSMA^high^IL-6^low^). RNAseq was performed and relative expression of genes (normalised expression) assessed using cufflinks (version 2.0.2) with default settings (25).

### Preparation of drugs, cell viability, cell cycle, cell death and senescence assays

Description is included in *Supplementary Materials and Methods*.

### Measurement of cystine uptake, glutathione synthesis, oxidative stress and glutamate efflux

Description is included in *Supplementary Materials and Methods*.

### 3D co-culture models, matrix contractility assays

Description is included in *Supplementary Materials and Methods*.

#### Transgenic pancreatic cancer mouse model and genetic ablation of SLC7A11

Genetically engineered mouse model (GEMM) experiments using the KC (Kras-mutated) and KPC (Kras- and p53-mutated) mouse model (29) [Alleles used: Pdx-1-promoter, lox-stop-lox-KrasG12D/+ allele, and lox-stop-lox-Trp53R172H/+ ± *Slc7a11^fl/fl^* (IMPC (MGI:1347355)] were genotyped by Transnetyx (Cordoba, TN, USA). *Pancreatic intraepithelial neoplasia (PanIN) scoring: Slc7a11^+/+^* KC (KC) and *Slc7a11^fl/fl^* (KC with conditional *Slc7a11* knockout under Pdx-1 promoter) mice were sampled at 70 days of age and PanINs scored from whole H&E sections and normalised to mm^2^ of section. *Survival:* KPC and KPC *Slc7a11^fl/fl^* mice were monitored at least 3 times weekly and sampled when exhibiting clinical signs of PDAC (abdominal swelling, jaundice, hunching, piloerection and weight loss). Slc7A11 knockout was confirmed by RNA in situ hybridisation and Western blot (**Figure S5**). RNA in situ hybridisation (ISH) was performed on formalin-fixed KPC tumour sections. RNA ISH (RNAscope) was performed according to the manufacturer’s protocol (ACD RNAscope 2.0 High Definition–Brown) for *Slc7a11* (Basecope probe targets floxed exon 3). Western blot was performed as above, except the following antibodies were used SLC7A11 (Cell Signaling Technology, Cat. 98051, 1:1000) and HSP90 (Cell Signaling Technology, Cat. 4875, 1:1000).

#### Star nanoparticle synthesis

Star nanoparticles (Star 3) were synthesised as previously described by our team (30). The purified core cross-linked star nanoparticle was analysed by GPC, NMR and FTIR after purification to determine its composition [final composition: f oligoethylene glycol methyl ether methacrylate (OEGMA)/f Dimethylaminoethyl methacrylate (DMEAMA) 14.5/85.5 mol %; Mn = 155,000 g/mol (± 5000g/mol); Average Size DLS = 28 (+/- 5nm); Average Zeta potential = 40 (+/-3)]. The nanoparticle was solubilised in methanol and dialysed with acidic water (pH = 3.0) for 24 h, and then further dialysed using water (pH = 6.5) for 48 h, then freeze-dried.

#### Orthotopic pancreatic cancer mouse model and SLC7A11 inhibition studies

Luciferase-expressing MiaPaCa-2 cells were established as described (43). These cells were co-implanted with human CAFs (10^6^ of each) into the tail of the pancreas of 8 week-old female BalbC nude mice, as described (43, 49). ***Star 3-siRNA gene silencing efficiency study:*** 6-weeks post-implant, mice were treated with 3mg/kg control-siRNA (antisense: 5’-GAACUUCAGGGUCAGCUUGCCG) or SLC7A11-siRNA (antisense: 5’-AGACCCAAUAAGUUUGCCG) complexed to Star 3 nanoparticles, intravenously once daily for three days. ***Star 3-siRNA therapeutic study:*** 4 weeks post-implant, mouse were randomised based on luminescence as described (43) (**Figure S7C**), then treated with 3mg/kg of control-siRNA or SLC7A11-siRNA complexed with Star 3, intravenously once daily for the first three days, followed by twice weekly for 4 weeks. Mice were co-treated intravenously with 10mg/kg Abraxane^®^ (10 mg/kg paclitaxel and 90mg/kg human albumin; Specialised Therapeutics Australia) or 90mg/kg human albumin (control), once weekly for 4 weeks. At end points, mice were humanely euthanised and organs/tumours harvested. Tumour volume was calculated by calliper measurement with operator blinded to treatment. Tumour fragments were 4% paraformaldehyde-fixed for histology, frozen in Tissue-Tek^®^ Optimal Cutting Temperature Compound (O.C.T; VWR International) for fluorescence analyses or snap frozen for protein extraction. Metastases were detected by ex vivo luminescent imaging (>600 counts) and confirmed by H&E as previously described (43).

### Measurement of collagen content and immunohistochemistry for SLC7A11, Alpha Smooth Muscle Actin (αSMA) and CD31

Full description of methods is in *Supplementary Materials and Methods*.

#### Statistical Analyses

Statistical comparisons were performed using two-tailed student t-test (2 groups) or ANOVA (≥3 groups; post-hoc tests: Dunn’s multiple comparison and Sidak’s multiple comparison). Analyses were performed using GraphPad Prism. Comparisons of univariate time to event (survival) were performed using the log-rank test and hazard ratios calculated from the Cox proportional hazards (PH) model. Multivariate associations between variables and time to event were contained from PH regression and survival curves calculated using the method of Kaplan-Meier (KM). Where tumour and stroma scores correlated with outcome, baseline variables associated with predicting scores were examined by multivariate logistic regression. Survival analyses were performed using Analysis of Censored and Correlated Data (ACCoRD) V6.4 Boffin. A p-value ≤0.05 was considered statistically significant.

#### Study Approval

All studies involving the use of human specimens and cell lines were approved by the UNSW Sydney human ethics committee (approvals: HC14039, HC180973, HREC13/023) and the German Technical University of Munich human ethics committee (approval: 5510/12). Animal studies were approved by the UNSW Sydney animal care and ethics committee (approval: ACEC 16/25B) for orthotopic mouse models and by local ethical review committee at University of Glasgow according to UK Home Office regulations (licence: 70/8646) for transgenic mouse models.

## DATA AVAILABILITY

All data generated or analysed during this study are included in this published article (and its Supplementary Information) and can be made available upon reasonable request. Expression array data for the PACA-AU cohort is publicly available through the ICGC data portal (*https://dcc.icgc.org/projects/PACA-AU*).

## AUTHOR CONTRIBUTIONS

GS, JM and AA share an equal author position. The order of equal authors was based on relative contribution of the co-first authors to experimental design, analysis of results and composition of the manuscript. GS, JM and PP designed and performed experiments, interpreted data, and wrote the manuscript. AA, CK, JY, RMCI, SN, JL, NR, JL, EG-A, JK performed *in vitro* and *in vivo* orthotopic tumour experiments. JH performed cystine uptake assays. JH and NT provided intellectual guidance on interpretation of metabolism-related results. CB and TPD synthesised and characterised Star 3 nanoparticles. ME isolated CAFs and MA isolated PSCs used in this study. PT, TRC, BAP, and JLC performed in vitro collagen plug assays and collagen analysis. SF, AKN, ADC, and OJS performed transgenic mouse experiments. DG, AC, AJ, AG and KH provided guidance on SLC7A11 scoring and analysis in the PDAC patient cohort study. VG performed statistical analyses for the PDAC patient cohort study. JM provided guidance on all *in vivo* aspects of the study. We also acknowledge the contribution of the Australian Pancreatic Cancer Genome Initiative in providing the valuable PDAC patient specimens and survival data utilised in the PDAC patient cohort study. **Australian Pancreatic Cancer Genome Initiative (APGI): Garvan Institute of Medical Research** Amber L. Johns^1^, Anthony J Gill^1, 5^, David K. Chang^1, 22^, Lorraine A. Chantrill^1,8^, Angela Chou^1,5^, Marina Pajic^1^, Angela Steinmann^1^, Mehreen Arshi^1^, Ali Drury^1^, Danielle Froio^1^, Ashleigh Parkin^1^, Paul Timpson^1^, David Hermann^1^. **QIMR Berghofer Medical Research Institute** Nicola Waddell^2^, John V. Pearson^2^, Ann-Marie Patch^2^, Katia Nones^2^, Felicity Newell^2^, Pamela Mukhopadhyay^2^, Venkateswar Addala^2^, Stephen Kazakoff^2^, Oliver Holmes^2^, Conrad Leonard^2^, Scott Wood^2^, Christina Xu^2^. **University of Melbourne, Centre for Cancer Research** Sean M. Grimmond^3^, Oliver Hofmann^3^. **University of QLD, IMB** Angelika Christ^4^, Tim Bruxner^4^. **Royal North Shore Hospital** Jaswinder S. Samra^5^, Jennifer Arena^5^, Nick Pavlakis^5,^ Hilda A. High^5^, Anubhav Mittal^5^. **Bankstown Hospital** Ray Asghari^6^, Neil D. Merrett^6^, Darren Pavey^6^, Amitabha Das^6^. **Liverpool Hospital** Peter H. Cosman^7^, Kasim Ismail^7^, Chelsie O’Connnor^7^. **St Vincent’s Hospital** Alina Stoita^8^, David Williams^8^, Allan Spigellman^8^**. Westmead Hospital** Vincent W. Lam^9,^ Duncan McLeod^9^, Adnan M. Nagrial^1,9^, Judy Kirk^9^. **Royal Prince Alfred Hospital, Chris O’Brien Lifehouse** James G. Kench^10^, Peter Grimison^10^, Caroline L. Cooper^10^, Charbel Sandroussi^10^, Annabel Goodwin^7,10^. **Prince of Wales Hospital** R. Scott Mead^1,11^, Katherine Tucker^11^, Lesley Andrews^11^. **Fremantle Hospital** Michael Texler^12^, Cindy Forest^12^, Krishna P. Epari^12^, Mo Ballal^12^, David R. Fletcher^12^, Sanjay Mukhedkar^12^. **St John of God Healthcare** Nikolajs Zeps^14^, Maria Beilin^14^, Kynan Feeney^14^. **Royal Adelaide Hospital** Nan Q Nguyen^15^, Andrew R. Ruszkiewicz^15^, Chris Worthley^15^. **Flinders Medical Centre** John Chen^16^, Mark E. Brooke-Smith^16^, Virginia Papangelis^16^. **Envoi Pathology** Andrew D. Clouston^17^, Patrick Martin^17^. **Princess Alexandria Hospital** Andrew P. Barbour^18^, Thomas J. O’Rourke^18^, Jonathan W. Fawcett^18^, Kellee Slater^18^, Michael Hatzifotis^18^, Peter Hodgkinson^18^. **Austin Hospital** Mehrdad Nikfarjam^19^. **Johns Hopkins Medical Institutes** James R. Eshleman^20^, Ralph H. Hruban^20^, Christopher L. Wolfgang^20^, Mary Hodgin^20^. **ARC-Net Centre for Applied Research on Cancer** Aldo Scarpa^21^, Rita T. Lawlor^21^, Stefania Beghelli^21^, Vincenzo Corbo^21^, Maria Scardoni^21^, Claudio Bassi^21^. **University of Glasgow** Andrew V Biankin^1, 22^, Judith Dixon^22^, Craig Nourse^22^, Nigel B. Jamieson^22^. ^1^The Kinghorn Cancer Centre, Garvan Institute of Medical Research, 370 Victoria Street, Darlinghurst, Sydney, New South Wales 2010, Australia. ^2^QIMR Berghofer Medical Research Institute, 300 Herston Rd, Herston, Queensland 4006, Australia. ^3^University of Melbourne, Centre for Cancer Research, Victorian Comprehensive Cancer Centre, 305 Grattan Street, Melbourne, Victoria 3000, Australia. ^4^ Institute for Molecular Bioscience, University of QLD, St Lucia, Queensland 4072, Australia. ^5^Royal North Shore Hospital, Westbourne Street, St Leonards, New South Wales 2065, Australia. ^6^Bankstown Hospital, Eldridge Road, Bankstown, New South Wales 2200, Australia. ^7^Liverpool Hospital, Elizabeth Street, Liverpool, New South Wales 2170, Australia. ^8^ St Vincent’s Hospital, 390 Victoria Street, Darlinghurst, New South Wales, 2010 Australia. ^9^Westmead Hospital, Hawkesbury and Darcy Roads, Westmead, New South Wales 2145, Australia. ^10^Royal Prince Alfred Hospital, Missenden Road, Camperdown, New South Wales 2050, Australia. ^11^Prince of Wales Hospital, Barker Street, Randwick, New South Wales 2031, Australia. ^12^Fremantle Hospital, Alma Street, Fremantle, Western Australia 6959, Australia. ^13^Sir Charles Gairdner Hospital, Hospital Avenue, Nedlands, Western Australia 6009, Australia. ^14^St John of God Healthcare, 12 Salvado Road, Subiaco, Western Australia 6008, Australia. ^15^Royal Adelaide Hospital, North Terrace, Adelaide, South Australia 5000, Australia. ^16^Flinders Medical Centre, Flinders Drive, Bedford Park, South Australia 5042, Australia. ^17^Envoi Pathology, 1/49 Butterfield Street, Herston, Queensland 4006, Australia. ^18^Princess Alexandria Hospital, Cornwall Street & Ipswich Road, Woolloongabba, Queensland 4102, Australia. ^19^Austin Hospital, 145 Studley Road, Heidelberg, Victoria 3084, Australia. ^20^Johns Hopkins Medical Institute, 600 North Wolfe Street, Baltimore, Maryland 21287, USA. ^21^ARC-NET Center for Applied Research on Cancer, University of Verona, Via dell’Artigliere, 19 37129 Verona, Province of Verona, Italy. ^22^Wolfson Wohl Cancer Research Centre, Institute of Cancer Sciences, University of Glasgow, Garscube Estate, Switchback Road, Bearsden, Glasgow, Scotland G61 1BD, United Kingdom). All authors have reviewed and approved this manuscript.

## MATERIALS AND CORRESPONDENCE

Please address all correspondence and request for materials to A/Prof Phoebe Phillips.

## ACKNOWLEDGEMENTS

We would like to thank Dr Andrea Nunez and Ms Amanda Mawson for their assistance in mouse monitoring. We would also like to acknowledge the Flow Cytometry, Biomedical Imaging, and Biological Resource Imaging Facilities within the Mark Wainwright Analytical Centre at the University of New South Wales for their technical support. Biospecimens and clinical data were provided by the Australian Pancreatic Cancer Genome Initiative (APGI, www.pancreaticcancer.net.au) which is supported by an Avner Pancreatic Cancer Foundation Grant, www.avnersfoundation.org.au. We would also like to acknowledge our collaborating community consumers Mr Gino Iori and Ms Claire Harvey for their invaluable input on the project and helping with consumer sections of grant applications. Funding: This research was made possible by major funding from NHMRC project grant (Phillips, McCarroll, Morton, Goldstein, APP1144108) and the Avner Foundation (Avner Innovation Grant [Phillips, McCarroll, Goldstein, Holst, Morton, Davis and Sharbeen, APCF0050618]; www.avnersfoundation.org.au). The following sources supported author contributions and research: NHRMC project grants (Cox and Chitty, APP1140125), NHMRC CDF-I (Phillips, APP1024896), NHMRC CDF-II (Cox, APP1158590), NHMRC Senior Research Fellowship (Timpson, APP1136974), Len Ainsworth Pancreatic Cancer Fellowship and support from Suttons, Cancer-Institute NSW ECF/CDFs (Phillips 08/ECF/1-37; Sharbeen, CDF181166; McCarroll, CDF102; Cox, CDF171105), Cancer Institute NSW Innovation Grant (Phillips, 09/RFG/2-58), Cancer Council NSW (Cox and Chitty, RG19-09), Cancer Australia (Phillips, McCarroll and Goldstein, APP1126736), Translational Cancer Research Network and Australian Postgraduate Award Scholarships (Akerman), Australian Government Research Training Program Scholarship & UNSW Sydney Scientia PhD Scholarship (Kokkinos), Cure Cancer Australia (Sharbeen, APP1122758), Cancer Research UK Core Funding and Grand Challenge grants (Sansom, Campbell, Najumudeen and Fey, A17196, A21139, A25045), Pancreatic Cancer UK Future Leaders Academy (Sansom, Fey). Competing Interests: The authors have no conflicts of interest to declare. Data and materials availability: All data related to this study can be found in the paper or the Supplementary Materials. More specific PDAC patient cohort data can be obtained through the Australian Pancreatic Cancer Genome Initiative. We are willing to share CAF cell lines, which would require appropriate human ethics approvals and an MTA with our laboratory.

## Reference

1. Siegel RL, Miller KD, and Jemal A. Cancer statistics, 2019. CA Cancer J Clin. 2019;69(1):7–34.

2. McCarroll JA, Naim S, Sharbeen G, Russia N, Lee J, Kavallaris M, et al. Role of pancreatic stellate cells in chemoresistance in pancreatic cancer. Front Physiol. 2014;5:141.

3. Pereira BA, Vennin C, Papanicolaou M, Chambers CR, Herrmann D, Morton JP, et al. CAF Subpopulations: A New Reservoir of Stromal Targets in Pancreatic Cancer. Trends Cancer. 2019;5(11):724–41.

4. Vaziri-Gohar A, Zarei M, Brody JR, and Winter JM. Metabolic Dependencies in Pancreatic Cancer. Front Oncol. 2018;8:617.

5. Sato H, Shiiya A, Kimata M, Maebara K, Tamba M, Sakakura Y, et al. Redox imbalance in cystine/glutamate transporter-deficient mice. The Journal of biological chemistry. 2005;280(45):37423–9.

6. Sato H, Tamba M, Ishii T, and Bannai S. Cloning and expression of a plasma membrane cystine/glutamate exchange transporter composed of two distinct proteins. The Journal of biological chemistry. 1999;274(17):11455–8.

7. Sato H, Tamba M, Kuriyama-Matsumura K, Okuno S, and Bannai S. Molecular cloning and expression of human xCT, the light chain of amino acid transport system xc. Antioxid Redox Signal. 2000;2(4):665–71.

8. Banjac A, Perisic T, Sato H, Seiler A, Bannai S, Weiss N, et al. The cystine/cysteine cycle: a redox cycle regulating susceptibility versus resistance to cell death. Oncogene. 2008;27(11):1618–28.

9. Wang K, Jiang J, Lei Y, Zhou S, Wei Y, and Huang C. Targeting Metabolic-Redox Circuits for Cancer Therapy. Trends Biochem Sci. 2019;44(5):401–14.

10. Chen RS, Song YM, Zhou ZY, Tong T, Li Y, Fu M, et al. Disruption of xCT inhibits cancer cell metastasis via the caveolin-1/beta-catenin pathway. Oncogene. 2009;28(4):599–609.

11. Gout PW, Buckley AR, Simms CR, and Bruchovsky N. Sulfasalazine, a potent suppressor of lymphoma growth by inhibition of the x(c)- cystine transporter: A new action for an old drug. Leukemia. 2001;15(10):1633–40.

12. Guan J, Lo M, Dockery P, Mahon S, Karp CM, Buckley AR, et al. The xc- cystine/glutamate antiporter as a potential therapeutic target for small-cell lung cancer: use of sulfasalazine. Cancer Chemother Pharmacol. 2009;64(3):463–72.

13. Huang Y, Dai Z, Barbacioru C, and Sadee W. Cystine-glutamate transporter SLC7A11 in cancer chemosensitivity and chemoresistance. Cancer Res. 2005;65(16):7446–54.

14. Ji X, Qian J, Rahman SMJ, Siska PJ, Zou Y, Harris BK, et al. xCT (SLC7A11)-mediated metabolic reprogramming promotes non-small cell lung cancer progression. Oncogene. 2018;37(36):5007–19.

15. Lyons SA, Chung WJ, Weaver AK, Ogunrinu T, and Sontheimer H. Autocrine glutamate signaling promotes glioma cell invasion. Cancer Res. 2007;67(19):9463–71.

16. Lo M, Ling V, Wang YZ, and Gout PW. The xc- cystine/glutamate antiporter: A mediator of pancreatic cancer growth with a role in drug resistance. British journal of cancer. 2008;99(3):464–72.

17. Lo M, Ling V, Low C, Wang YZ, and Gout PW. Potential use of the anti-inflammatory drug, sulfasalazine, for targeted therapy of pancreatic cancer. Current oncology. 2010;17(3):9–16.

18. Arensman MD, Yang XS, Leahy DM, Toral-Barza L, Mileski M, Rosfjord EC, et al. Cystine-glutamate antiporter xCT deficiency suppresses tumor growth while preserving antitumor immunity. Proc Natl Acad Sci U S A. 2019;116(19):9533–42.

19. Daher B, Parks SK, Durivault J, Cormerais Y, Baidarjad H, Tambutte E, et al. Genetic ablation of the cystine transporter xCT in PDAC cells inhibits mTORC1, growth, survival and tumor formation via nutrient and oxidative stresses. Cancer Res. 2019.

20. Ohman KA, Hashim YM, Vangveravong S, Nywening TM, Cullinan DR, Goedegebuure SP, et al. Conjugation to the sigma-2 ligand SV119 overcomes uptake blockade and converts dm-Erastin into a potent pancreatic cancer therapeutic. Oncotarget. 2016;7(23):33529–41.

21. Badgley MA, Kremer DM, Maurer HC, DelGiorno KE, Lee HJ, Purohit V, et al. Cysteine depletion induces pancreatic tumor ferroptosis in mice. Science. 2020;368(6486):85–9.

22. Kshattry S, Saha A, Gries P, Tiziani S, Stone E, Georgiou G, et al. Enzyme-mediated depletion of l-cyst(e)ine synergizes with thioredoxin reductase inhibition for suppression of pancreatic tumor growth. NPJ Precis Oncol. 2019;3:16.

23. Sousa CM, Biancur DE, Wang X, Halbrook CJ, Sherman MH, Zhang L, et al. Pancreatic stellate cells support tumour metabolism through autophagic alanine secretion. Nature. 2016;536(7617):479–83.

24. Zhang W, Trachootham D, Liu J, Chen G, Pelicano H, Garcia-Prieto C, et al. Stromal control of cystine metabolism promotes cancer cell survival in chronic lymphocytic leukaemia. Nat Cell Biol. 2012;14(3):276–86.

25. Ohlund D, Handly-Santana A, Biffi G, Elyada E, Almeida AS, Ponz-Sarvise M, et al. Distinct populations of inflammatory fibroblasts and myofibroblasts in pancreatic cancer. J Exp Med. 2017;214(3):579–96.

26. Bailey P, Chang DK, Nones K, Johns AL, Patch AM, Gingras MC, et al. Genomic analyses identify molecular subtypes of pancreatic cancer. Nature. 2016;531(7592):47–52.

27. Ishii T, Bannai S, and Sugita Y. Mechanism of growth stimulation of L1210 cells by 2- mercaptoethanol in vitro. Role of the mixed disulfide of 2-mercaptoethanol and cysteine. The Journal of biological chemistry. 1981;256(23):12387–92.

28. Zhu S, Zhang Q, Sun X, Zeh HJ, 3rd, Lotze MT, Kang R, et al. HSPA5 Regulates Ferroptotic Cell Death in Cancer Cells. Cancer Res. 2017;77(8):2064–77.

29. Hingorani SR, Wang L, Multani AS, Combs C, Deramaudt TB, Hruban RH, et al. Trp53R172H and KrasG12D cooperate to promote chromosomal instability and widely metastatic pancreatic ductal adenocarcinoma in mice. Cancer Cell. 2005;7(5):469–83.

30. Teo J, McCarroll JA, Boyer C, Youkhana J, Sagnella SM, Duong HT, et al. A Rationally Optimized Nanoparticle System for the Delivery of RNA Interference Therapeutics into Pancreatic Tumors in Vivo. Biomacromolecules. 2016;17(7):2337–51.

31. Ma Z, Zhang H, Lian M, Yue C, Dong G, Jin Y, et al. SLC7A11, a component of cysteine/glutamate transporter, is a novel biomarker for the diagnosis and prognosis in laryngeal squamous cell carcinoma. Oncol Rep. 2017;38(5):3019–29.

32. Zhang L, Huang Y, Ling J, Zhuo W, Yu Z, Luo Y, et al. Overexpression of SLC7A11: a novel oncogene and an indicator of unfavorable prognosis for liver carcinoma. Future Oncol. 2018;14(10):927–36.

33. Maurer C, Holmstrom SR, He J, Laise P, Su T, Ahmed A, et al. Experimental microdissection enables functional harmonisation of pancreatic cancer subtypes. Gut. 2019;68(6):1034–43.

34. Yang Q, Li K, Huang X, Zhao C, Mei Y, Li X, et al. lncRNA SLC7A11-AS1 Promotes Chemoresistance by Blocking SCF(beta-TRCP)-Mediated Degradation of NRF2 in Pancreatic Cancer. Mol Ther Nucleic Acids. 2020;19:974–85.

35. Koppula P, Zhang Y, Shi J, Li W, and Gan B. The glutamate/cystine antiporter SLC7A11/xCT enhances cancer cell dependency on glucose by exporting glutamate. The Journal of biological chemistry. 2017;292(34):14240–9.

36. Shin CS, Mishra P, Watrous JD, Carelli V, D’Aurelio M, Jain M, et al. The glutamate/cystine xCT antiporter antagonizes glutamine metabolism and reduces nutrient flexibility. Nat Commun. 2017;8:15074.

37. Alonso V, Linares V, Belles M, Albina ML, Sirvent JJ, Domingo JL, et al. Sulfasalazine induced oxidative stress: a possible mechanism of male infertility. Reprod Toxicol. 2009;27(1):35–40.

38. Wahl C, Liptay S, Adler G, and Schmid RM. Sulfasalazine: a potent and specific inhibitor of nuclear factor kappa B. J Clin Invest. 1998;101(5):1163–74.

39. Weber CK, Liptay S, Wirth T, Adler G, and Schmid RM. Suppression of NF-kappaB activity by sulfasalazine is mediated by direct inhibition of IkappaB kinases alpha and beta. Gastroenterology. 2000;119(5):1209–18.

40. Elahi-Gedwillo KY, Carlson M, Zettervall J, and Provenzano PP. Antifibrotic Therapy Disrupts Stromal Barriers and Modulates the Immune Landscape in Pancreatic Ductal Adenocarcinoma. Cancer Res. 2019;79(2):372–86.

41. Ene-Obong A, Clear AJ, Watt J, Wang J, Fatah R, Riches JC, et al. Activated pancreatic stellate cells sequester CD8+ T cells to reduce their infiltration of the juxtatumoral compartment of pancreatic ductal adenocarcinoma. Gastroenterology. 2013;145(5):1121–32.

42. Miller BW, Morton JP, Pinese M, Saturno G, Jamieson NB, McGhee E, et al. Targeting the LOX/hypoxia axis reverses many of the features that make pancreatic cancer deadly: inhibition of LOX abrogates metastasis and enhances drug efficacy. EMBO Mol Med. 2015;7(8):1063–76.

43. McCarroll JA, Sharbeen G, Liu J, Youkhana J, Goldstein D, McCarthy N, et al. betaIII- tubulin: a novel mediator of chemoresistance and metastases in pancreatic cancer. Oncotarget. 2015;6(4):2235–49.

44. Sharbeen G, McCarroll J, Liu J, Youkhana J, Limbri LF, Biankin AV, et al. Delineating the Role of betaIV-Tubulins in Pancreatic Cancer: betaIVb-Tubulin Inhibition Sensitizes Pancreatic Cancer Cells to Vinca Alkaloids. Neoplasia. 2016;18(12):753–64.

45. Sharbeen G, Youkhana J, Mawson A, McCarroll J, Nunez A, Biankin A, et al. MutY- Homolog (MYH) inhibition reduces pancreatic cancer cell growth and increases chemosensitivity. Oncotarget. 2017;8(6):9216–29.

46. Ouyang H, Mou L, Luk C, Liu N, Karaskova J, Squire J, et al. Immortal human pancreatic duct epithelial cell lines with near normal genotype and phenotype. Am J Pathol. 2000;157(5):1623–31.

47. Apte MV, Haber PS, Applegate TL, Norton ID, McCaughan GW, Korsten MA, et al. Periacinar stellate shaped cells in rat pancreas: identification, isolation, and culture. Gut. 1998;43(1):128–33.

48. Bachem MG, Schneider E, Gross H, Weidenbach H, Schmid RM, Menke A, et al. Identification, culture, and characterization of pancreatic stellate cells in rats and humans. Gastroenterology. 1998;115(2):421–32.

49. Vonlaufen A, Joshi S, Qu C, Phillips PA, Xu Z, Parker NR, et al. Pancreatic stellate cells: Partners in crime with pancreatic cancer cells. Cancer Res. 2008;68(7):2085–93.

50. Morton JP, Timpson P, Karim SA, Ridgway RA, Athineos D, Doyle B, et al. Mutant p53 drives metastasis and overcomes growth arrest/senescence in pancreatic cancer. Proc Natl Acad Sci U S A. 2010;107(1):246–51.

51. Edfors F, Hober A, Linderback K, Maddalo G, Azimi A, Sivertsson A, et al. Enhanced validation of antibodies for research applications. Nat Commun. 2018;9(1):4130.

52. Wang Q, Hardie RA, Hoy AJ, van Geldermalsen M, Gao D, Fazli L, et al. Targeting ASCT2-mediated glutamine uptake blocks prostate cancer growth and tumour development. J Pathol. 2015;236(3):278–89.

53. Conway JRW, Vennin C, Cazet AS, Herrmann D, Murphy KJ, Warren SC, et al. Three-dimensional organotypic matrices from alternative collagen sources as pre-clinical models for cell biology. Sci Rep. 2017;7(1):16887.

54. Vennin C, Chin VT, Warren SC, Lucas MC, Herrmann D, Magenau A, et al. Transient tissue priming via ROCK inhibition uncouples pancreatic cancer progression, sensitivity to chemotherapy, and metastasis. Sci Transl Med. 2017;9(384).

55. Vennin C, Melenec P, Rouet R, Nobis M, Cazet AS, Murphy KJ, et al. CAF hierarchy driven by pancreatic cancer cell p53-status creates a pro-metastatic and chemoresistant environment via perlecan. Nat Commun. 2019;10(1):3637.

56. Cazet AS, Hui MN, Elsworth BL, Wu SZ, Roden D, Chan CL, et al. Targeting stromal remodeling and cancer stem cell plasticity overcomes chemoresistance in triple negative breast cancer. Nat Commun. 2018;9(1):2897.

57. Cicchi R, Kapsokalyvas D, De Giorgi V, Maio V, Van Wiechen A, Massi D, et al. Scoring of collagen organization in healthy and diseased human dermis by multiphoton microscopy. J Biophotonics. 2010;3(1-2):34–43.

58. Mayorca-Guiliani AE, Madsen CD, Cox TR, Horton ER, Venning FA, and Erler JT. ISDoT: in situ decellularization of tissues for high-resolution imaging and proteomic analysis of native extracellular matrix. Nat Med. 2017;23(7):890–8.

