## Supplementary methods, figures and tables for "Cancer-associated fibroblasts in pancreatic ductal adenocarcinoma determine response to SLC7A11 inhibition"

### SUPPLEMENTARY ITEMS

1. Supplementary materials and methods
2. Supplementary figures
3. Supplementary tables

### 1. SUPPLEMENTARY METHODS

##### Quantitative real-time PCR (qPCR).

Quantitative real-time PCR (qPCR) was performed using Power SYBR® green kit (ThermoFisher Scientific) run on an Applied Biosystems ViiA7 real-time PCR platform. Primers were obtained from Qiagen: SLC7A11 (Cat. QT00002674); 18S ribosomal RNA (housekeeper gene for standardisation of PSC vs CAF SLC7A11 and SLC3A2 qPCR analysis, Cat. QT00199367); SLC3A2 (Cat. QT00085897); $\beta_2$-microglobulin (housekeeping gene for standardisation of knockdown qPCR analysis, Cat. QT00199367). Concentrations were calculated based on ΔCt standard curves and standardised to $\beta_2$-microglobulin.

##### Western blot for SLC7A11.

Protein lysates were prepared using Cell Lysis Buffer (Cell Signalling Technologies), quantified by BCA Assay, electrophoresed on a 10% SDS-PAGE gel and transferred to nitrocellulose membranes as described (43), except samples were not boiled prior to electrophoresis. Membranes were probed with the following antibodies: α-tubulin (Housekeeping protein; Sigma Aldrich, Cat. T9026, 1:1000), SLC7A11 (for human: Cell Signaling Technologies, Cat. 12691, Abcam, Cat. ab37185,1:1000; for mouse: Cell Signaling Technologies Cat. 98051, 1:1000), then anti-rabbit or anti-mouse secondary antibodies (DAKO, Cat. P0447, P0448). Blots were visualised using Amersham Western Blotting Detection Reagents (GE Healthcare Life Sciences) scanned using LAS4000 scanner and quantified using ImageQuant TL (GE Healthcare Life Sciences).

##### siRNA transfection.

Human and mouse CAFs and PDAC cells were transfected with 100nM smartpool On-target plus human SLC7A11-siRNA (Dharmacon, Cat. L-007612-01), human SLC7A11-siRNA single sequence siRNA [Dharmacon, Cat. LU-007612-01 (part of the set-of-4 SLC7A11 ON-TARGET siRNA, J-007612-01)], mouse SLC7A11-siRNA (Dharmacon, Cat. L-047420-01) or control non-silencing siRNA (ns-siRNA; Dharmacon, Cat. D-001810-01-50)] using Lipofectamine 2000 (ThermoFisher Scientific) as described (43-45).

**Preparation of drugs.**

Sulfasalazine (SSZ, Sigma Aldrich, Cat. S0883-10G) was dissolved in 0.1M NaOH, 1x phosphate buffered saline (PBS), pH7.5 and passed through a 0.22μm filter. Erastin (Cayman Chemical, Cat. 17754) was reconstituted in sterile dimethyl sulfoxide (DMSO; Sigma Aldrich). Ferrostatin-1 (Sigma-Aldrich, Cat. SML0583) was dissolved in DMSO. N-acetyl-L-cysteine (NAC; Sigma-Aldrich, Cat. A9165) was dissolved in MilliQ $H_2O$ and filter-sterilised and passed through a 0.22μm filter.

**Cell viability assays.**

Cell proliferation or viability was measured by trypan blue exclusion and cell counting kit 8 (CCK8) assays as described (43-45) at the following treatment times: (i) 72h post-transfection with siRNA; (ii) 48h post-treatment with 0.2-0.4mM SSZ; (iii) 24-48h post-treatment with 40μM erastin; (iv) 72h post-seeding of shRNA stable lines. *Co-treatments:* (i) 0-60μM tert-butyl hydroperoxide (tBHP; oxidative stress inducer; Sigma Aldrich, Cat. 458139) was used in the final 24h of assays; (ii) 0-66μM 2-mercaptoethanol (2-ME) was used in parallel with SSZ; (iii) 2μM Ferrostatin was used in the final 24h of assays.

**Cell cycle, cell death and senescence assays.**

**Ferroptosis assay:** Adhered CAFs (i) 72h post-transfection with siRNA, (ii) 48h post-seeding of stable shRNA CAFs, (iii) 9h post-treatment with 40 μM erastin or 0.2% DMSO (control) were assayed for ferroptosis using a colorimetric assay (Glutathione Peroxidase Assay Kit; Abcam, Cat. ab102530) according to the manufacturer's instructions. **Cell cycle assay:** Cell cycle analysis of CAFs was performed by propidium iodide (PI) staining and flow cytometry on a BD Fortessa, 72h post-transfection, as previously described (45). **Cell senescence assay:** Senescence was assessed in CAFs 72h post-transfection with ns-siRNA or SLC7A11-siRNA or 72h post-treatment with SSZ. The induction of senescence in CAFs was measured using a Senescence β-galactosidase Cell Staining Kit (Cell Signalling Technologies, Cat. 9860) according to the manufacturer's instructions. **Measurement of autophagy:** Autophagy was assessed in CAFs 72h post-transfection with ns-siRNA or SLC7A11-siRNA. Western blot was performed for the microtubule-associated protein Light Chain 3 isoform B (LC3B), to check for conversion from cytosolic LC3BI to autophagosome-bound LC3BII. Western blot was performed as per

SLC7A11 Western blot procedure, except samples were boiled at 95°C for 5min prior to loading. GAPDH was used as a loading control. Western blots were probed with the following antibodies: LC3B (D11) XP® (1:1000; Cell Signaling Technology, Cat. 3868S); GAPDH (1:50,000, Housekeeper protein; Abcam, Cat. ab8245). The blots were scanned using LAS4000 scanner and quantified using ImageQuant TL software (GE Healthcare). **Apoptosis assay:** CAFs were transfected with SLC7A11-siRNA or ns-siRNA then treated 48h post-transfection ± tBHP, for a further 24h. Adherent and floating cells were harvested and total apoptosis detected using an AnnexinV-PE/7AAD kit (BD Bioscience, Cat. 559763) according to the manufacturer's instructions and analysed on a BD LSRFortessa$^{TM}$ flow cytometer.

**Measurement of cystine uptake.**

Non-transfected CAFs (SSZ experiments) or CAFs, 48h post-transfection with siRNA, were seeded at 15,000 cells/well into Wallac isoplate 96–well plates. The following day, cystine uptake was assayed immediately (knockdown experiments) or 2-3h post-treatment with SSZ (SSZ maintained during assay). Cystine uptake was measured using a previously described assay (52). 0.1μCi [14C]–L-cystine (Perkin Elmer) was added to cell mixture in Hanks Buffered Salt Solution and transferred to a humidified incubator for 15min at 37$^{o}$C. Cells were then lysed with 1% Triton-X100 in PBS and 180μl scintillant added per well. Radioactivity was measured using a liquid scintillation counter (Perkin Elmer).

**Assessing glutathione synthesis, oxidative stress and glutamate efflux.**

**Measurement of intracellular glutathione:** Total intracellular glutathione was measured in whole cell lysates (prepared: (i) 72h post-transfection with siRNA; (ii) 72h post-seeding of shRNA stable lines; (iii) after 48h 0.25mM SSZ; (iv) 16h post-0.25mM SSZ with or without 1mM NAC) using ApoGSH$^{TM}$ Glutathione Colormetric Assay (BioVision, Cat. K261-100) or Glutathione Assay Kit (Cayman Chemical, 703002) according to manufacturer's instructions. **Detection of intracellular ROS using CellROX:** CAFs were incubated in 0.4mM SSZ or transfected with ns-siRNA or SLC7A11-siRNA. Seventy-two hours post-transfection or after addition of SSZ, CAFs were incubated with 325μM tert-butyl hydroperoxide (tBHP) for 1h. CAFs were then stained with CellROX green reagent (Life Technologies, C10444) according to the manufacturer's instructions. Fluorescence was analysed on a BD LSRFortessa$^{TM}$ flow cytometer. **Glutamate efflux assay:** CAFs transfected 72h prior with ns-siRNA or SLC7A11-siRNA, were washed and incubated in M2 buffer (Sigma Aldrich) for 4h at 37°C in a humidified atmosphere. Glutamate was measured in neat cell supernatant using an Amplex Red Glutamic Acid/Glutamate Oxidase Assay Kit (Thermofisher Scientific; Cat. A12221) according to the manufacturer's instructions. Fluorescence was measured using an EnSpire Multimode Plate Reader (PerkinElmer) at 571/586nm (excitation/emission) wavelengths.

**3D co-culture models.**

MiaPaCa-2 PDAC cells and CAFs were transfected with ns-siRNA or SLC7A11-siRNA, then the following assays performed. **Spheroid outgrowth assay:** After 24h, 2500 of each cell type was co-seeded in suspension. Spheroids were allowed to form over 24h under normal culture conditions, then transferred to soft-agarose medium (20% FBS, 4% L-glutamine, 0.33% agarose in low glucose DMEM:F12 medium) and overlayed on to 50µL pre-set soft-agarose medium (0.5% agarose) in a 96-well round-bottom plate. Outgrowth was quantified 6 days later from brightfield images using ImageJ. **Spheroid growth assay:** MiaPaCa-2 PDAC cells and CAFs stably expressing scramble-shRNA or SLC7A11-shRNA seq 1 (see method above) were seeded into low adherence round-bottom 96-well plates at a ratio of 1:3 (2500 MiaPaCa-2 + 7500 CAFs) in 100µl complete medium. After 24h culture to allow spheroids to form, supernatant on spheroids was replaced with a 1:1 mix of complete medium: Corning® Matrigel® Growth Factor Reduced (GFR) Basement Membrane Matrix (In Vitro Technologies, Cat. 354230). Spheroids were allowed to grow for 7 days under normal culture conditions and daily bright field photos taken on an inverted light microscope.

**Matrix contractility assay.**

Organotypic assays were adapted from published protocols (53, 54) (supplementary methods). Rat-tail tendon collagen was extracted with 0.5 mM acetic acid to a concentration of 2.5 mg/ml. 0.3-1x$10^5$ CAFs (n=4) were embedded in 1.25 mL of rat-tail collagen I. Once polymerised, fibroblast-collagen matrices were allowed to contract in phenol red free IMDM containing 10% FBS, 4 mM GlutaMAX and 1% penicillin/streptomycin for six days with images taken at Day 2, 4 and 6. Collagen plug area was quantified from brightfield images. At Day 6, collagen matrices were fixed in 10% neutral buffered formalin, then paraffin embedded and sectioned for collagen analysis as described (53-55). Second Harmonic Generation (SHG) signal was acquired using a Leica DMI 6000 SP8 inverted confocal multiphoton microscope with a 25x 0.95 NA water objective, as previously described. Four representative regions of interest (512x512) were imaged per sample, with a line average of 4 and over a 3D z-stack (100 μm depth with a z-step size of 2.52 μm). SHG signal intensity was measured using Matlab (MathWorks), as previously described (54, 55). Paraffin-embedded samples were cut into 4μm sections and stained with 0.1% picrosirius red (Polysciences) for fibrillar collagen, according to manufacturer's instructions. Picrosirius signal intensity was measured using Matlab (MathWorks), as previously described (54, 55). Polarised light imaging was performed as previously described (54, 55). Collagen fibre network organisation was characterised using grey-level co-occurrence matrix (GLCM) analysis as previously described (54, 55). The correlation curves represent the similarity in signal intensity between pixels acquired via single-plane SHG imaging of collagen fibres (line average for SHG acquisition: 16). GLCM analysis was performed in Matlab (Mathworks) as previously described (54, 55).

**Measurement of collagen content and organisation in pancreatic tumour sections (orthotopic and transgenic PDAC models).**

***Picrosirius red staining and measurement of total collagen content and fibril density (polarised light analysis):*** Pancreatic tumour tissue was fixed in 4% paraformaldehyde and embedded in paraffin. Paraffin-embedded samples were cut into 5μm sections and stained with 0.1% picrosirius red for fibrillar collagen, performed through the UNSW Mark Wainwright Analytical Centre Biomedical Imaging Facility (UNSW Sydney, Australia). Sections were counter-stained with methyl green. ***Total collagen content:*** Collagen content was calculated from representative images per tumour at 20x magnification (excluding necrotic regions and tumour edges, average tumour coverage = 13%) using the ImageJ colour deconvolution module followed by measurement of percent of area positive for picrosirius red per ROI. These were then averaged for each tumour. ***Polarised light analysis:*** This analysis was performed on the same ROIs as those used for the total collagen content analysis. As previously described (53-55), polarised light imaging was performed on a Leica DM6000 fitted with a polariser in combination with a transmitted light analyser. Quantitative intensity measurements of fibrillar collagen content and birefringent signal were carried out using in house scripts in ImageJ. For each polarised light image, Hue-Saturation-Balance (HSB) thresholding was applied, where $0 \geq H \leq 27$ | $0 \geq S \leq 255$ | $5 \geq B \leq 255$ was used for red-orange (high birefringent) fibres, $28 \geq H \leq 47$ | $0 \geq S \leq 255$ | $5 \geq B \leq 255$ for yellow (medium birefringent) fibres, and $48 \geq H \leq 140$ | $0 \geq S \leq 255$ | $5 \geq B \leq 255$ for green (low birefringent) fibres. The relative area (as a % of total fibres [$0 \geq H \leq 140$ | $0 \geq S \leq 255$ | $5 \geq B \leq 255$]) was then calculated. ***Second harmonics analysis of collagen organisation and directionality*****:** Collagen was visualised in OCT-embedded tumour sections (15 μm thick) by Second Harmonics Generation (SHG) confocal microscopy. SHG signal was acquired using a HC PL FLUOTAR 20.0x0.50 dry objective on an inverted Leica SP5 confocal microscope. Excitation source was a Mai Tai DeepSee multiphoton laser, operating at 200Hz and tuned to a wavelength of 840 nm (line average = 5). Intensity was recorded with Non-Descanned Detectors (NDD). For each sample, 3 representative regions of interest of 775 μm x 775 μm were imaged over a 3D z-stack (15 μm depth with a z-step size of 1.5 μm). Maximum intensity projections of these z-stacks were used for grey-level co-occurrence matrix (GLCM) and orientation analyses. ***Grey-level co-occurrence matrix (GLCM) Analysis (Fibril Organisation):*** Fibrillar collagen organisation was characterised using grey-level co-occurrence matrix (GLCM) analysis as previously described (54, 56). GLCM provides a readout of the texture of a sample by quantifying the similarity between pixels across an image (57). GLCM analysis was performed in MATLAB (Mathworks) as previously described (57). The GLCM correlation parameter for each image was calculated using looped operation of the plug-in for 0°, 90°, 180° and 270° directions. Normalised correlations were calculated in MATLAB (MathWorks) and the mean correlation was plotted against the distance in GraphPad software. ***Collagen orientation analysis.*** Collagen orientation analysis was carried out as previously described (56, 58). In brief, fibre orientation analysis was performed on SHG images using an in-house ImageJ macro where structure tensors were derived from the local orientation and isotropic properties of pixels that make up collagen fibrils. Within each input image, these tensors were evaluated for each pixel by computing the continuous spatial derivatives in the x and y dimensions using a cubic B-spline interpolation. From this, the local predominant orientation was obtained. The peak alignment (measured in degrees) of fibres was then determined, and the frequency of fibre alignment calculated.

**Immunohistochemistry for SLC7A11, Alpha Smooth Muscle Actin (αSMA) and CD31.** Immunohistochemistry was performed in paraformaldehyde-fixed and paraffin-embedded tumour sections, as described (43-45) using the following antibodies: SLC7A11 (Cell Signaling Technologies, Cat. 12691; 1:100), αSMA (Orthotopic: Sigma, Cat. A5228, 1:5000; KPC: Sigma, Cat. A2547, 1:25000), CD31 (Taylor Bio-Medical, Cat. DIA-310, 1:50), Rat IgG isotype control (Abcam, Cat. ab18703), rabbit IgG isotype control (DAKO, Cat. X0903), biotinylated anti-rabbit secondary antibody (Vector Laboratories, Cat. BA-1000; 1:400), biotinylated anti-rat secondary antibody (Vector laboratories, Cat. BA-4000; 1:100) and Vectastain® ABC kit (Vector laboratories). SLC7A11 staining intensity, αSMA area coverage and CD31-positive open and closed blood vessels were analysed in representative images using colour deconvolution in ImageJ. SLC7A11 staining intensity was calculated from three representative images per tumour (average area coverage = 10% per section, excluding necrotic regions) using the ImageJ colour deconvolution module followed by measurement of pixel intensity. Pixel intensity was converted to optical density (intensity of staining) using the following formula: log(maximum intensity of pixels/average intensity of pixels). αSMA area coverage was calculated from representative images per tumour at 20x magnification (excluding necrotic regions; average tumour coverage = 21%). Following colour deconvolution in ImageJ, DAB staining was quantified as a fraction of the total area per region of interest, then averaged for each tumour. CD31-positive open and closed blood vessels were manually counted in representative tumour regions (excluding necrotic regions; average tumour coverage = 55%). Observers were blinded to treatment groups in all above analyses.

### 2. SUPPLEMENTARY FIGURES

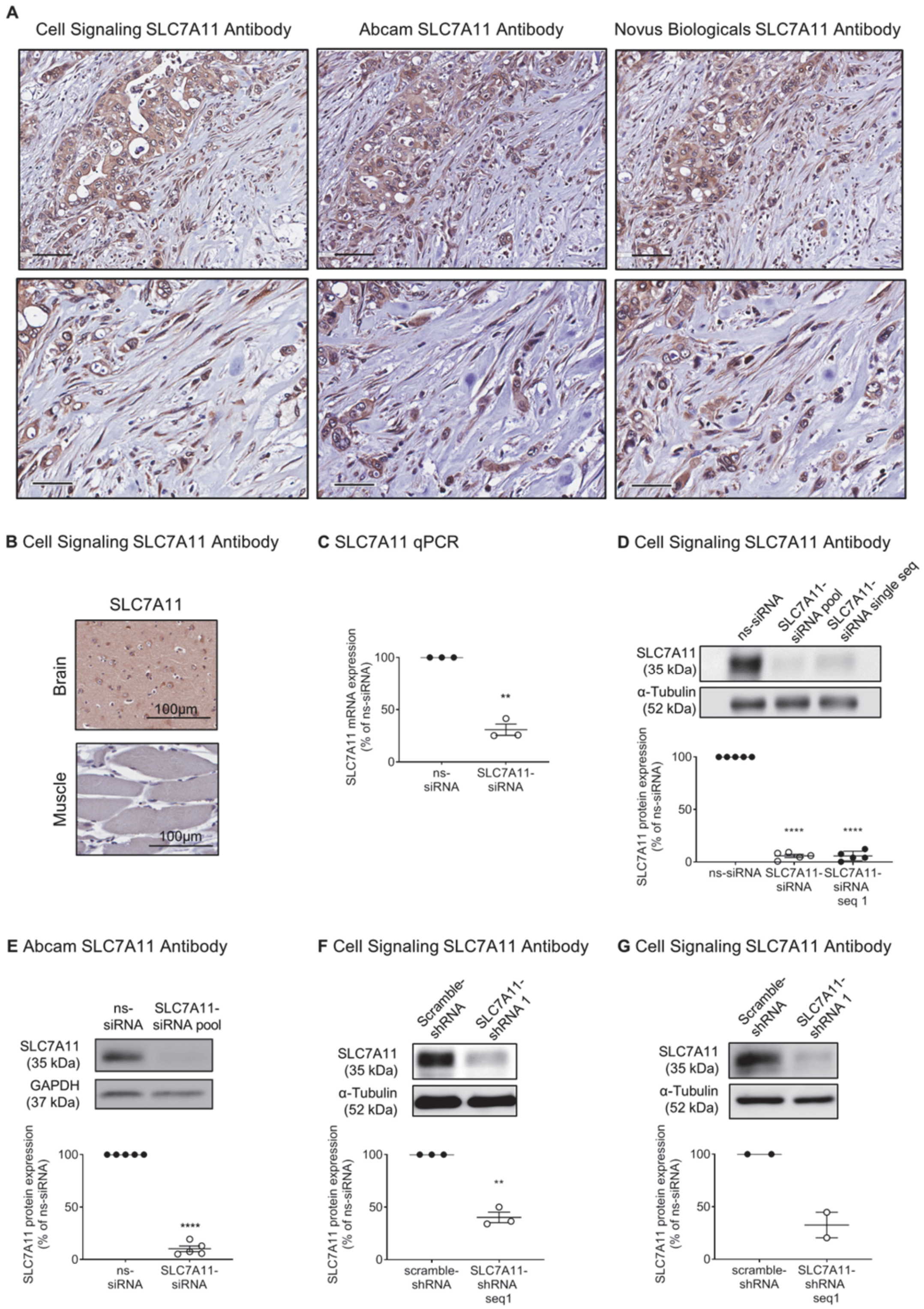

**Figure S1: Validation of SLC7A11 antibodies and SLC7A11 knockdown in CAFs and PDAC cells *in vitro*.** A) Comparison of immunohistochemistry for SLC7A11 in APGI ICGC PDAC patient tumour tissue using 3 different antibody clones from Cell Signaling, Abcam and Novus Biologicals. B) Positive (brain) and negative (skin) controls for SLC7A11 IHC in the APGI ICGC cohort. C) Quantitative real-time PCR analysis of SLC7A11 silencing in total RNA extracts from CAFs, 72h post-transfection with ns-siRNA or SLC7A11-siRNA. Circles indicate replicates, lines indicate mean±s.e.m., asterisks indicate significance (****p≤0.0001, n=3; student t-test). D-E) Representative Western blot [using (D) Cell signaling or (E) Abcam antibodies] and densitometry of SLC7A11 in total protein extracts from CAFs, 72h after transfection with control siRNA (ns-siRNA), SLC7A11-siRNA pool or SLC7A11-siRNA single sequence (SLC7A11-siRNA single seq). α-tubulin or GAPDH were used as loading controls. Circles indicate independent CAF cell lines, lines indicate mean±s.e.m., asterisks indicate significance (****p≤0.0001, n=5; D= One-way ANOVA, E=student t-test). F-G) As per (D) except total protein extracts from CAFs and MiaPaCa-2 PDAC cells stably expressing scramble-shRNA or SLC7A11-shRNA sequence 1 (SLC7A11-shRNA seq 1) were used (**p≤0.01; student t-test).

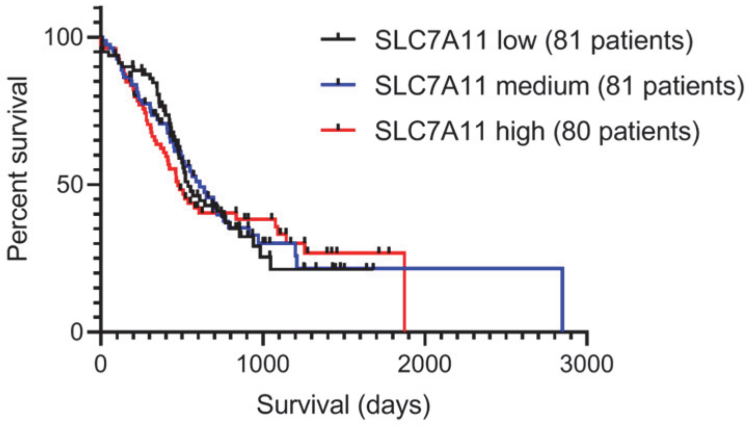

**Figure S2: SLC7A11 mRNA expression in the ICGC PDAC cohort does not predict patient survival.** Expression array data for SLC7A11 was analysed in PDAC patients from the APGI ICGC cohort. Patients were broken into tertiles based on SLC7A11 expression (low, medium, high) and correlated with overall patient survival. Kaplan-Meier survival curves are shown. Curves were not significantly different.

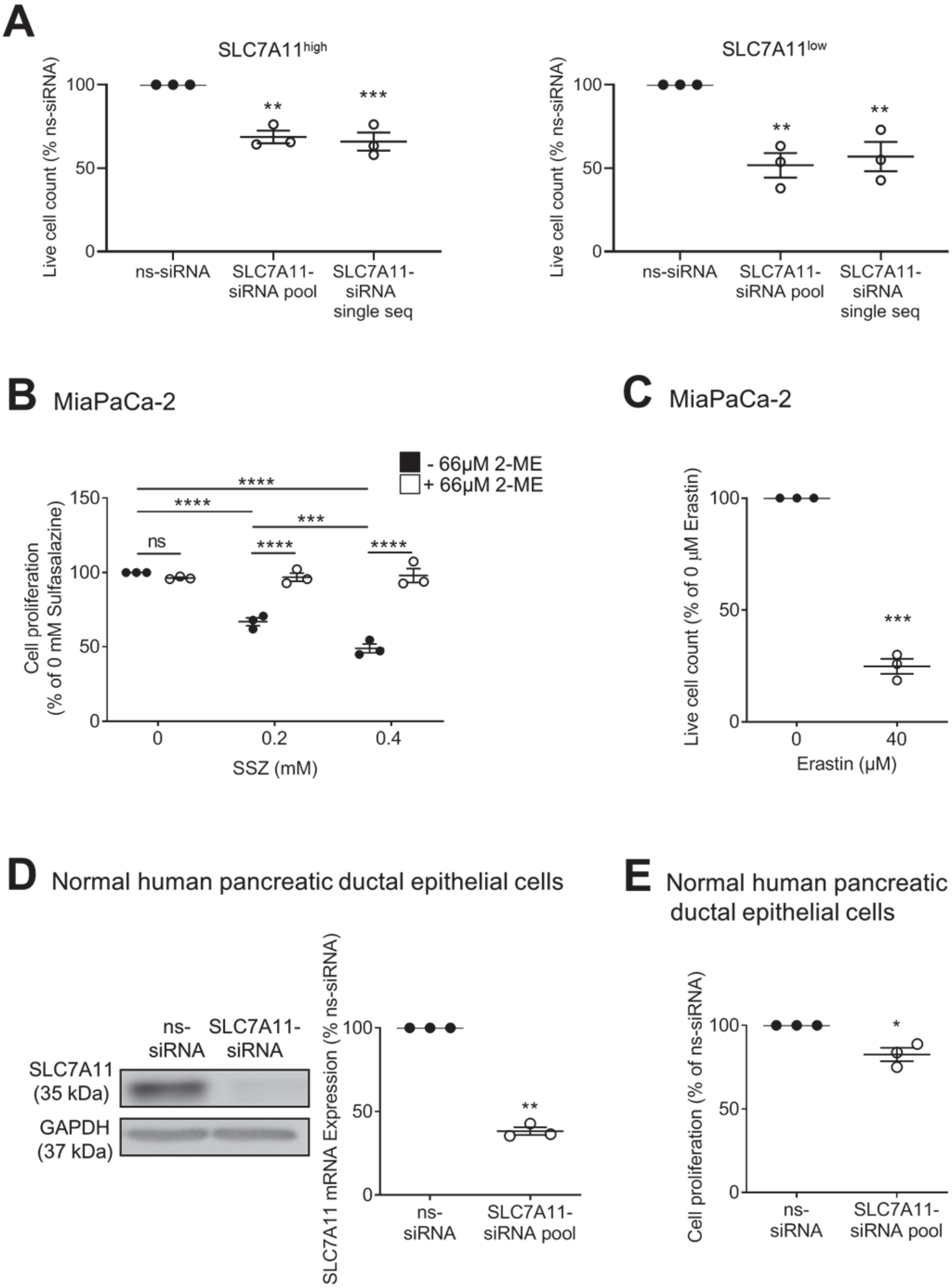
A
SLC7A11high
SLC7A11low
Live cell count (% ns-siRNA)
100
50
0
**
***
**
**
ns-siRNA
SLC7A11-siRNA pool
SLC7A11-siRNA single seq
B MiaPaCa-2
- 66μM 2-ME
+ 66μM 2-ME
Cell proliferation (% of 0 mM Sulfasalazine)
150
100
50
0
ns
****
***
SSZ (mM)
0
0.2
0.4
C MiaPaCa-2
Live cell count (% of 0 μM Erastin)
***
Erastin (μM)
40
D Normal human pancreatic ductal epithelial cells
ns-siRNA
SLC7A11-siRNA
SLC7A11 (35 kDa)
GAPDH (37 kDa)
SLC7A11 mRNA Expression (% ns-siRNA)
**
ns-siRNA
SLC7A11-siRNA pool
E Normal human pancreatic ductal epithelial cells
Cell proliferation (% of ns-siRNA)
*
ns-siRNA
SLC7A11-siRNA pool

**Figure S3: Anti-proliferative effect of SLC7A11 knockdown in CAFs and the effect of SLC7A11 inhibition in MiaPaCa-2 PDAC cells and normal human pancreatic ductal epithelial (HPDE) cells.** A) Live cell counts (trypan blue exclusion) of CAFs expressing low (CAF line 6 in Figure 1B) or high basal levels of SLC7A11 (CAF line 1 in Figure 1B), 72h post-transfection with control siRNA (ns-siRNA), SLC7A11-siRNA pool or SLC7A11-siRNA single sequence (SLC7A11-siRNA single seq). Circles indicate replicate experiments (n=3). B) Cell proliferation (cell counting kit 8 absorbance) of MiaPaCa-2 PDAC cells treated with sulfasalazine (SSZ) ± 66μM 2-mercaptoethanol (2-ME), as a % of controls. Circles indicate replicates (n=3). C) Live cell counts (trypan blue exclusion) of MiaPaCa-2 cells treated with erastin, as a fraction of controls. Circles indicate replicates (n=3). D) Representative Western blot and qPCR for SLC7A11 knockdown in HPDE cells, 72h post-transfection. Circles indicate replicates (n=3). E) Cell proliferation (cell counting kit 8 absorbance) of HPDE cells 72h post-transfection with ns-siRNA or SLC7A11-siRNA pool, as a % of controls. Circles indicate replicates (n=3). One-way ANOVA used for (A-B), student t-test used for (C-E). Asterisks in all graphs indicate significance (ns = not significant, *p≤0.05, **p≤0.01, ***p≤0.001, ****p≤0.0001). Replicate numbers for all CAF experiments refer to independent transfections/treatments using CAF cells isolated from different PDAC patients. Replicate numbers for all HPDE and MiaPaCa-2 experiments refer to independent transfections and treatments.

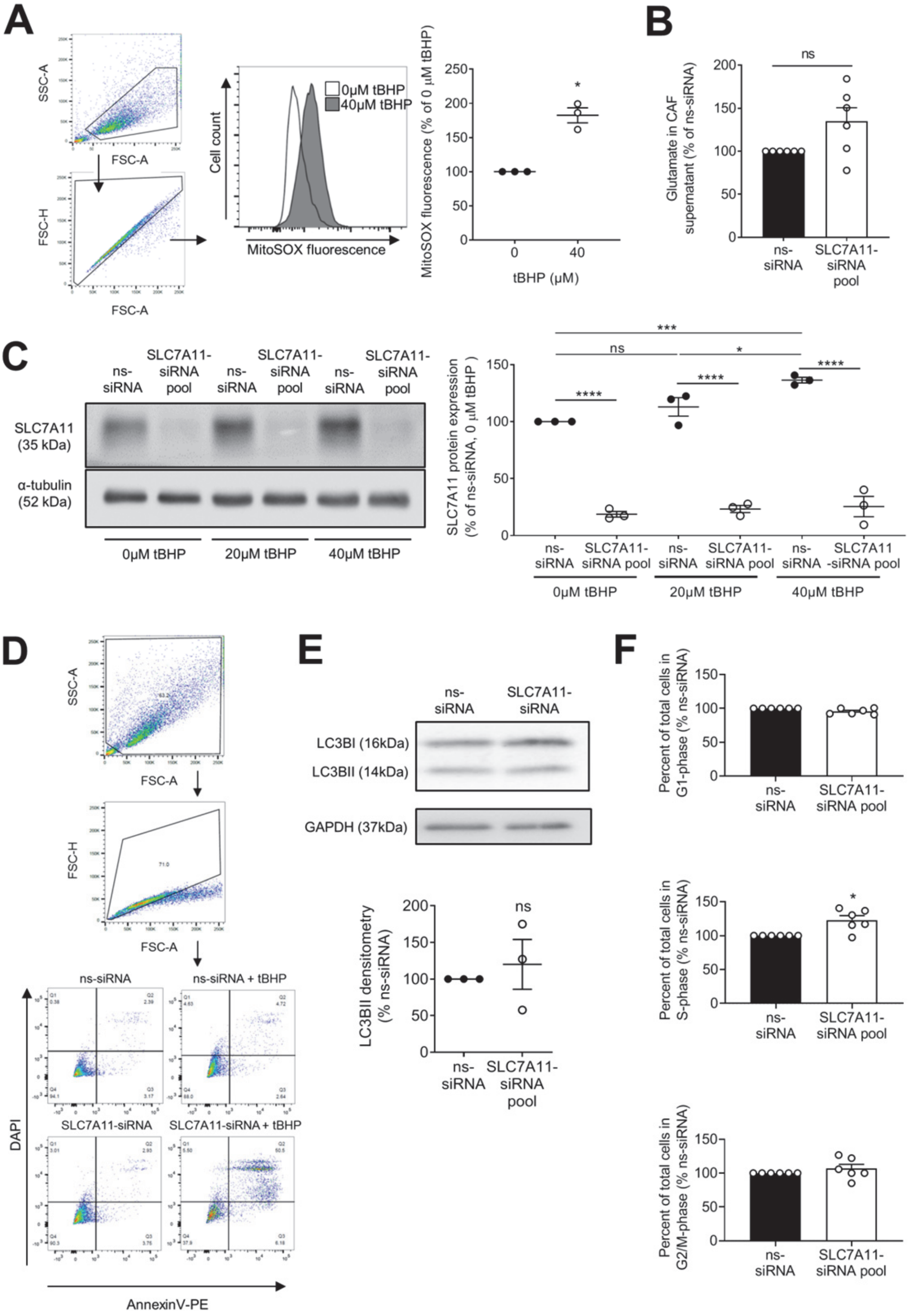
A
SSC-A
FSC-A
FSC-H
FSC-A
Cell count
MitoSOX fluorescence
0μM tBHP
40μM tBHP
MitoSOX fluorescence (% of 0 μM tBHP)
250
200
150
100
50
0
*
0
40
tBHP (μM)
B
ns
Glutamate in CAF supernatant (% of ns-siRNA)
200
150
100
50
0
ns-siRNA
SLC7A11-siRNA pool
C
ns-siRNA
SLC7A11-siRNA pool
ns-siRNA
SLC7A11-siRNA pool
ns-siRNA
SLC7A11-siRNA pool
SLC7A11 (35 kDa)
α-tubulin (52 kDa)
0μM tBHP
20μM tBHP
40μM tBHP
***
ns
*
****
****
****
SLC7A11 protein expression (% of ns-siRNA, 0 μM tBHP )
150
100
50
0
ns-siRNA
SLC7A11-siRNA pool
ns-siRNA
SLC7A11-siRNA pool
ns-siRNA
SLC7A11 -siRNA pool
0μM tBHP
20μM tBHP
40μM tBHP
D
SSC-A
FSC-A
FSC-H
FSC-A
ns-siRNA
ns-siRNA + tBHP
SLC7A11-siRNA
SLC7A11-siRNA + tBHP
DAPI
AnnexinV-PE
E
ns-siRNA
SLC7A11-siRNA
LC3BI (16kDa)
LC3BII (14kDa)
GAPDH (37kDa)
LC3BII densitometry (% ns-siRNA)
200
150
100
50
0
ns
ns-siRNA
SLC7A11-siRNA pool
F
Percent of total cells in G1-phase (% ns-siRNA)
150
100
50
0
ns-siRNA
SLC7A11-siRNA pool
Percent of total cells in S-phase (% ns-siRNA)
150
100
50
0
*
ns-siRNA
SLC7A11-siRNA pool
Percent of total cells in G2/M-phase (% ns-siRNA)
150
100
50
0
ns-siRNA
SLC7A11-siRNA pool

**Figure S4: SLC7A11 knockdown in PDAC CAFs does not affect glutamate secretion and is maintained in the presence of oxidative stress.** A) Gating strategy, representative histogram and quantification of the geometric mean of fluorescence peaks for CAFs treated with or without 40μM tBHP. Circles indicate replicates (n=3). B) Bars show average secreted glutamate in cell supernatants from CAFs transfected with ns-siRNA or SLC7A11-siRNA pool (as a fraction of control-siRNA), as assessed by colorimetric assay. Circles indicate replicates (n=6). C) Representative Western blot and densitometry of SLC7A11 in total protein extracts from CAFs 72h after transfection with ns-siRNA or SLC7A11-siRNA pool, and 48h post treatment with tBHP. α-tubulin = loading control. Circles indicate replicates (n=3). D) Gating strategy and representative AnnexinV-PE/DAPI flow cytometry pseudocolour plots for CAFs treated with ns-siRNA or SLC7A11-siRNA, with or without 40μM tBHP. E) Representative Western blot for LC3BI/II and densitometry for LC3BII (increases during autophagy) in total protein extracts from CAFs transfected with ns-siRNA or SLC7A11-siRNA. GAPDH = loading control. Circles indicate replicates (n=3). F) Bars represent fraction of total live cells in each cell cycle phase, as determined by DAPI stain and flow cytometry 72h after transfection with ns-siRNA or SLC7A11-siRNA pool (n=6). Bars/lines in all graphs = mean±s.e.m. Student t-test used for (A-B, E-F), one-way ANOVA used for (C). Asterisks in all graphs indicate significance (ns=not significant, *p≤0.05, ***p≤0.001, ****p≤0.0001). Replicate numbers for all CAF experiments refer to independent transfections/treatments using CAF cells isolated from different PDAC patients.

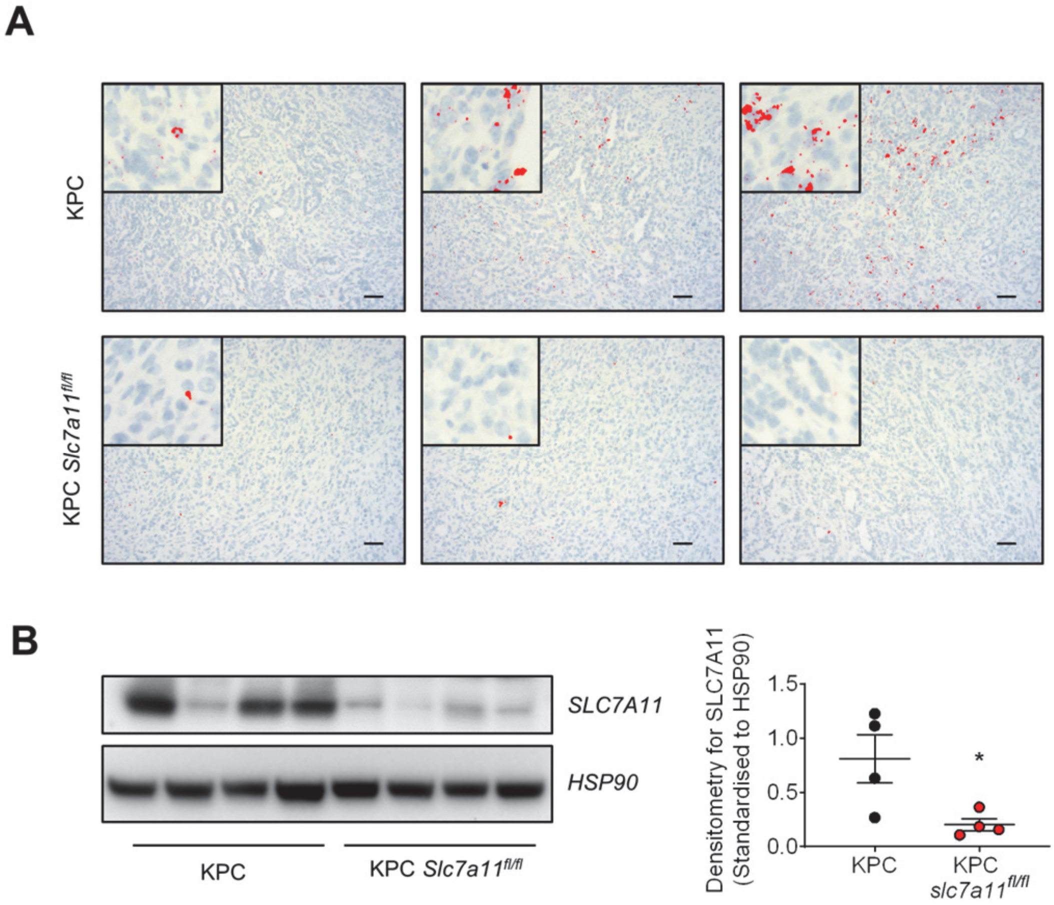

**Figure S5: Confirmation of SLC7A11 KO in the KPC transgenic model.** A) In situ hybridisation for SLC7A11 transcripts in KPC and KPC *slc7a11*$^{fl/fl}$ tumour sections. Red = SLC7A11 signal. B) Western blot for SLC7A11 in protein extracts from KPC and KPC *slc7a11*$^{fl/fl}$ mice. Graph shows densitometry for SLC7A11, standardised to HSP90. Dots represent individual mice, lines show mean±s.e.m. (* $p<0.05$, n=4; student t-test). Scale bars = 20µm.

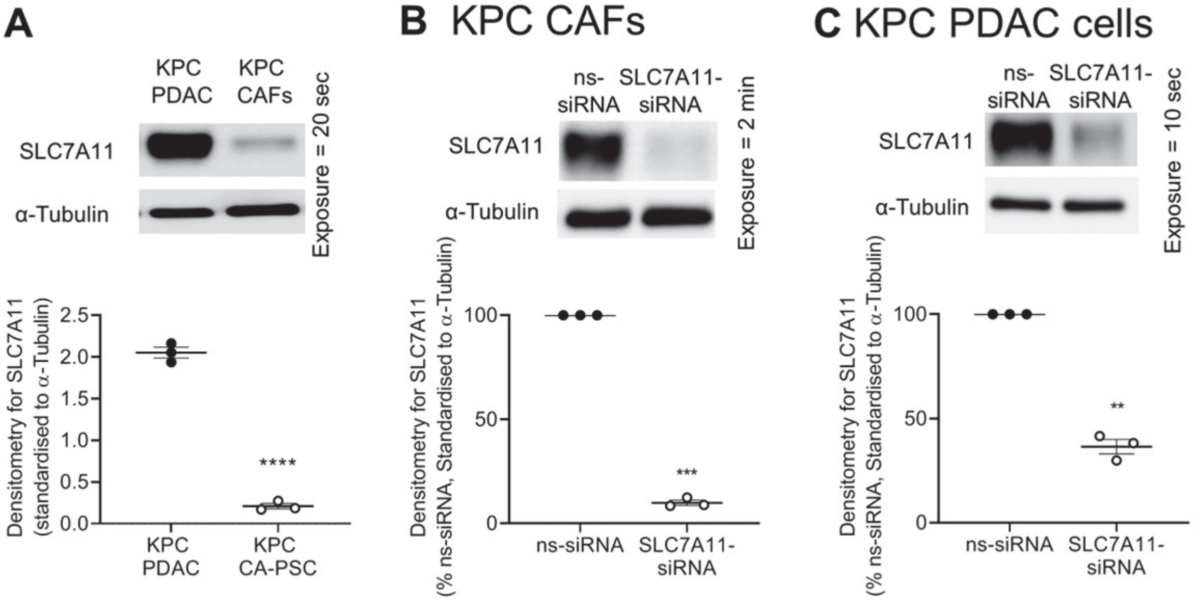

**Figure S6: Relative expression and knockdown of SLC7A11 in KPC PDAC cells and KPC CAFs**. A) Western blot comparing SLC7A11 in KPC PDAC cells and KPC CAFs. B) Western blot and densitometry for SLC7A11 knockdown in KPC CAFs (standardised to α-tubulin). C) As per B, expect KPC PDAC cell extracts were used. Exposure times for luminescence acquisition are shown next to each blot. Circles in all graphs indicate replicate experiments, lines in all graphs show mean±s.e.m. Student t-test used for all graphs, asterisks indicate significance (**p≤0.01,***p≤0.001,****p≤0.0001; n=3).

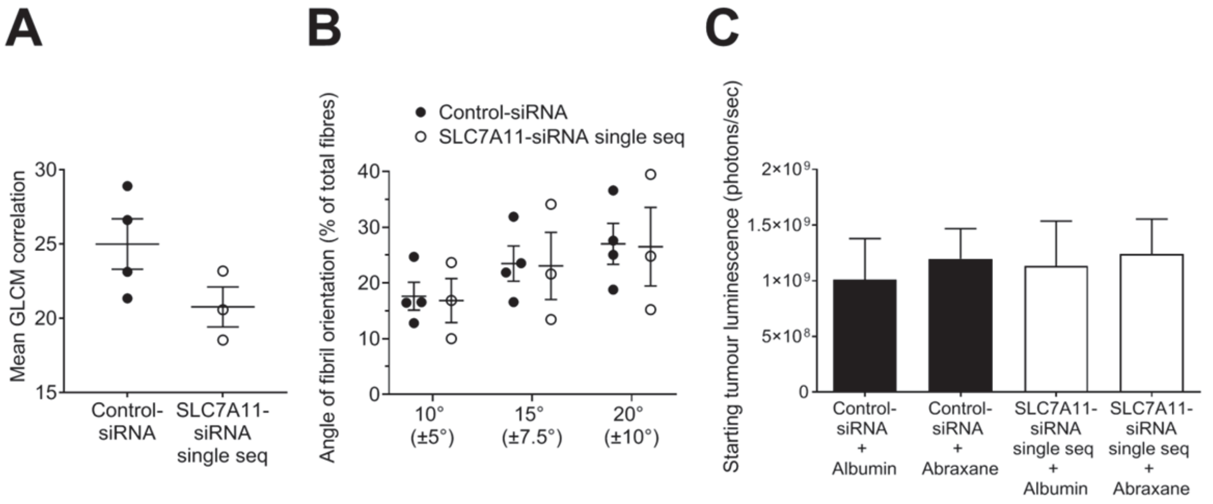

**Figure S7: Orthotopic model standardisation and collagen analysis.** A-B) Analysis of collagen fibril directionality and organisation in tumour sections at therapeutic model endpoint. Collagen was detected by second harmonics generation and the maximum intensity projections of 3 representative regions were used per tumour section. A) Grey-level co-occurrence matrix (GLCM) analysis. Graph shows mean correlation per group. Circles indicate individual mice, lines indicate mean±s.e.m. B) Graph shows the % of total fibrils with angles of ± $10^o$, $15^o$ and $20^o$ deflection from the centre axis. Circles indicate individual mice, lines indicate mean±s.e.m. C) Graph shows mean tumour luminescence (+s.e.m.) post-randomisation for therapeutic model of SLC7A11 knockdown (n=7-8 per group).

### 3. SUPPLEMENTARY TABLES

| | TA010 (myCAF 1) | TA011 (myCAF 2) | TA012 (quiescent PSC 1) | TA013 (quiescent PSC 2) | TA014 (iCAF 1) | TA015 (iCAF 2) | TA016 (iCAF 3) | TA017 (iCAF 4) |
|---|---|---|---|---|---|---|---|---|
| **Slc7a11 normalised expression** | 439.4 | 834.1 | 408.6 | 379.3 | 1122.0 | 1307.1 | 862.1 | 896.4 |
| | | **myCAF** | | **qPSC** | | | | **iCAF** |
| **Slc7a11 average normalised expression** | | 636.8 | | 394.0 | | | | 1046.9 |

**Supplementary Table 1: SLC7A11 is upregulated in iCAFS and myCAFs relative to normal pancreatic fibroblasts.** Summary of normalised SLC7A11 expression data from Ohlund et al (25) demonstrating higher average expression of SLC7A11 in immune-modulatory CAFs (iCAFs) and myofibrolast CAFs (myCAFs) relative to quiescent pancreatic fibroblasts (quiescent PSCs). Each column represents an independent culture of myCAFs, iCAFs or quiescent fibroblasts (refer to (25)).

| **Univariate analysis of parameters used in multivariate analysis** | | | | |
|---|---|---|---|---|
| **Parameter** | **HR** | **95%** | **CI** | **p-value** |
| Tumour score | 0.837 | 0.587 | 1.193 | 0.325 |
| Stroma score | 1.34 | 0.946 | 1.899 | 0.099 |
| Gender | 0.788 | 0.555 | 1.119 | 0.182 |
| Age at Diagnosis (years) | 0.851 | 0.425 | 1.705 | 0.649 |
| Smoker | 0.998 | 0.701 | 1.421 | 0.989 |
| Alcohol consumption | 0.93 | 0.646 | 1.338 | 0.695 |
| Margin Status | 1.721 | 1.19 | 2.487 | 0.004 |
| Lymph Nodes Involved | 1.409 | 0.908 | 2.184 | 0.126 |
| Perineural Invasion | 1.214 | 0.739 | 1.997 | 0.444 |
| Vascular Invasion | 1.982 | 1.336 | 2.941 | 0.001 |
| Overall Stage AJCC | 1.66 | 1.064 | 2.59 | 0.026 |
| Macroscopic Tumour Location | 0.885 | 0.458 | 1.709 | 0.716 |
| **Multivariate results (best subsets)** | **HR** | **95%** | **CI** | **p-value** |
| Stroma score | 1.45 | 1.016 | 2.07 | 0.041 |
| Vascular Invasion | 2.066 | 1.388 | 3.076 | 0.000 |

**Supplementary Table 2: SLC7A11 multivariate survival analysis parameters**

| Age at diagnosis | Number of patients |
|---|---|
| ≥50 | 144 |
| <50 | 11 |
| **Gender** | |
| Male | 85 |
| Female | 70 |
| **Ethnicity** | |
| Asian | 12 |
| Asian, White/Caucasian | 1 |
| Black/African | 1 |
| Pacific Islander | 1 |
| White/Caucasian | 140 |
| **Smoker** | |
| Ever | 81 |
| Never | 70 |
| Not reported | 4 |
| **Alcohol consumption** | |
| Ever | 88 |
| Never | 61 |
| Not reported | 6 |
| **Margin Status** | |
| R0 | 105 |
| R1 | 47 |
| R2 | 3 |
| **Macroscopic tumour location** | |
| Ampulla | 2 |
| Body | 11 |
| Head | 121 |
| Head (Uncinate) | 7 |
| Tail | 14 |

| Overall Stage | Number of patients |
|---|---|
| IA | 2 |
| IB | 2 |
| IIA | 34 |
| IIB | 113 |
| III | 0 |
| IV | 4 |
| **TNM Staging** | |
| T1 | 2 |
| T2 | 5 |
| T3 | 148 |
| N0 | 30 |
| N1 | 77 |
| N1a | 7 |
| N1b | 32 |
| NX | 1 |
| M0 | 7 |
| M1 | 4 |
| MX | 144 |
| **Perineural invasion** | |
| Yes | 131 |
| No | 23 |
| Not reported | 1 |
| **Vascular invasion** | |
| Yes | 95 |
| No | 56 |
| Not reported | 4 |
| **Recurrence at liver** | |
| Yes | 54 |
| No | 63 |
| No recurrence | 38 |

**Supplementary Table 3: APGI ICGC Patient Cohort Characteristics**
